# Exclusion from DNA double-strand breaks is a pervasive feature of the hnRNP family

**DOI:** 10.64898/2026.11.02.763712

**Authors:** Estelle Vincendeau, Leonie von Ondarza, Julie Klaric, Nikhil Venkatesh, Carmine Ceraolo, Stephanie Panier

## Abstract

Heterogeneous nuclear ribonucleoproteins (hnRNPs) are abundant RNA-binding proteins that have been linked individually to the DNA double-strand break (DSB) response, but their behaviour at damaged chromatin has only been examined one protein at a time and under disparate conditions. Here, we compare the core hnRNP family side by side by laser microirradiation and live-cell imaging. We find that 13 of 14 members are excluded from damaged chromatin, 10 directly and 3 after a transient phase of recruitment, establishing exclusion rather than accumulation as the predominant behaviour of this family. We characterised this behaviour in detail for hnRNPH1, which has not previously been linked directly to the DSB response. Its exclusion is sustained for several hours and also occurs at nuclease-induced breaks at a defined genomic locus. Importantly, exclusion is independent of the signalling pathways that drive protein recruitment to breaks and of transcriptional elongation, and instead requires the GYR low-complexity domain, with RNA binding contributing redundantly through three quasi-RNA recognition motifs. Finally, tethering hnRNPH1 to breaks through the RNF8 FHA domain selectively impairs 53BP1 accumulation without affecting γ-H2AX, MDC1 or RPA, and hnRNPH1 is excluded from 53BP1 nuclear bodies in unchallenged cells. Together, these findings establish protein exclusion as an active component of the DSB response and show that the presence of an hnRNP at damaged chromatin can determine which repair factors assemble at DNA break sites.

## INTRODUCTION

DNA double-strand breaks (DSBs) are the most dangerous form of DNA damage as they disrupt the physical continuity of the chromosomes. If left unrepaired or repaired inaccurately, they can cause chromosomal translocations, deletions and chromosome loss, and contribute to diseases such as immunodeficiency, neurodegeneration and cancer (1). To protect genome stability, cells counter this threat with a complex signalling network that detects DSBs, propagates their signals across the surrounding chromatin and determines how they are repaired (2). PARP1, a poly(ADP) ribose polymerase, is among the first factors at the DNA lesion, where it adds PAR chains both to itself and to neighbouring chromatin substrates to seed the early recruitment of DNA damage response factors (3–6). In parallel, the MRE11-RAD50-NBS1 (MRN) sensor complex also assembles at DSBs and activates the ATM kinase, which phosphorylates histone variant H2AX at S139 (also called γ-H2AX) over a megabase domain flanking the break site (7–11). γ-H2AX nucleates the assembly of the transducer protein MDC1, which in turn recruits the RNF8-RNF168 ubiquitylation cascade (12–16). RNF168-mediated ubiquitylation of histone H2A on K15 then licenses the recruitment of 53BP1 and BRCA1, whose mutual antagonism governs the extent of DNA end-resection at the break site and thereby DSB repair pathway choice (17–20). Indeed, together with its effectors RIF1 and the shieldin complex, 53BP1 limits DNA end-resection and channels repair towards non-homologous end-joining (NHEJ), which ligates broken DNA ends directly throughout interphase (18, 19, 21–30). In contrast, BRCA1, in complex with BARD1, displaces 53BP1 from damaged chromatin and, through its phospho-dependent interaction with CtIP, stimulates the endonucleolytic activity of the MRN complex to initiate short-range resection (18, 19, 31–33). Subsequent long-range resection generates the long 3’ single-stranded overhangs on which homologous recombination (HR)-directed DSB repair depends (34). Because HR uses the sister chromatid as a repair template, its activity is restricted to the S and G2 phases of the cell cycle.

In the course of DNA damage signalling and repair, the DSB response generates transient, dynamic compartments that concentrate signalling and repair factors locally around the break site (35–39). However, recruitment is not the only change in composition that these compartments undergo. For example, the poly (ADP)-ribose synthesised within seconds of DNA damage induction is likewise turned over within minutes to release the factors it recruited (6, 40, 41). In addition, transcription is silenced *in cis* across the chromatin domain flanking the break, and elongating RNA polymerase II is displaced and targeted for proteasomal degradation (42–44). The composition of damaged chromatin is therefore shaped in both directions, by recruitment and release, and on comparable timescales.

RNA metabolism is among the processes most immediately affected by these events. The transcriptional response at DSB sites is itself complex (45). It engages in multiple pathways rather than a single silencing mechanism, and, additionally, damage-induced transcripts have been implicated in the assembly of repair factors such as 53BP1 (46–48). Proteins that bind RNA are therefore exposed to dynamic changes in substrates and chromatin environment, and a growing number of RNA-binding proteins (RBPs) act locally at DSBs in ways unrelated to their canonical roles in gene expression (49).

Among RBPs, the heterogeneous ribonucleoprotein (hnRNP) family is most extensively connected to the chromatin-based DSB response (49). The family comprises 16 core members (named A to U) that are all non-enzymatic RNA-binding proteins best characterised for their functions in RNA synthesis, splicing, stability, transport and translation (50). They share a modular architecture of one, but more often, several RNA-binding domains (RBDs) and low-complexity domains (LCDs), which mediate protein-protein interactions, drive liquid-liquid phase separation, and function in cellular signalling and organisation. Several hnRNP members have been implicated in the DSB response, although their role is not always clear. For example, hnRNPUL1 and hnRNPUL2 promote DNA end-resection and HR-directed repair, while SAF-A (hnRNPU) has been linked to NHEJ (51, 52). FUS (hnRNPP2), RBMX (hnRNPG), CIRBP (hnRNPA18) and PCBP1 (hnRNPE1) have been linked to efficient DSB signalling and repair (53–59). In addition, many more hnRNPs can be found in the interactomes of central components of the DSB response such as MDC1, 53BP1, and BRCA1, indicating a broader involvement of the hnRNP family in DSB signalling or repair than is currently established (26, 60). Characterisation of hnRNPs at DSBs has largely been limited to those members that accumulate at damaged chromatin such as FUS, RBMX, hnRNPUL1 and hnRNPUL2 (52, 53, 55, 59). However, accumulation is not always sustained. Both FUS and SAF-A are recruited to breaks in a PAR-dependent manner and subsequently excluded from damaged chromatin (51, 58). In addition, hnRNPA2B1 and PCBP1 have also been reported to be excluded from laser microirradiation-induced DNA breaks, with hnRNPA2B1 release repressing local transcription (57, 61). Neither the prevalence of exclusion across the hnRNP family nor its consequences for DNA repair factor assembly have so far been addressed.

Here, we show that exclusion from damaged chromatin is a pervasive feature of the hnRNP family. We found that 13 of 14 family members tested were excluded from laser microirradiation tracks, 10 directly and 3 after a transient phase of recruitment. We characterise this behaviour in detail for hnRNPH1, which had not been previously linked directly to the DSB response. hnRNPH1 exclusion is sustained for several hours and occurs independently of the DNA damage signalling events that drive protein recruitment at break sites. Instead, exclusion depends on the GYR low-complexity domain, which mediates phase separation *in vitro*, whereas RNA binding likely contributes redundantly through the protein’s three quasi-RNA recognition motifs (qRRMs). Finally, to test the consequences of retaining hnRNPH1 at damaged chromatin, we tethered it to DSBs by fusing it to the RNF8 FHA domain. Retained hnRNPH1 selectively impairs 53BP1 accumulation while leaving γ-H2AX, MDC1 and RPA recruitment intact, showing that the presence of an hnRNP at damaged chromatin can determine which repair factors assemble at the DNA lesion, and suggesting that their exclusion contributes to normal focus assembly.

## RESULTS

### Exclusion from DSBs is a pervasive feature of the hnRNP family

Despite evidence that the hnRNP family responds to DNA damage, hnRNPs have so far been examined at DSB sites one at a time, in different cell lines and under different microirradiation conditions (51–58). We therefore set out to compare the behaviour of hnRNP family members directly and under identical conditions. To this end, we carried out a targeted microscopy screen to determine the subcellular localisation of hnRNP family members following DSB induction (Figure 1 and Supplementary Figure 1). HeLa cells expressing GFP-tagged hnRNPs were subjected to laser microirradiation and followed by time-lapse imaging for 15 minutes to assess the spatiotemporal distribution of each protein at damaged chromatin. For each protein, relative fluorescence intensity along the irradiated line was quantified in at least 30 cells from three independent experiments, with GFP alone as a negative control. As a positive control, we used the minimal focus-forming region of 53BP1 (53BP1^MFFR^), which accumulated along the irradiated line within 5 minutes as previously reported (Supplementary Figure 1A) (62, 63). Of the 14 family members we examined, 13 were excluded from damaged chromatin. Ten hnRNPs (hnRNPH1, hnRNPD, PCBP1, hnRNPK, hnRNPL, hnRNPR, hnRNPA2B1, hnRNPF, PTBP1 (hnRNPI) and hnRNPQ) were excluded directly following DSB induction (Figure 1A-F and Supplementary Figure 1B-E). Exclusion was rapid and detectable within 30 to 60 seconds of damage induction. Three hnRNPs (FUS, RBMX and SAF-A) showed a biphasic behaviour, accumulating at damaged chromatin within 2 minutes as previously described (51, 53, 56, 58, 59, 64) before being excluded with exclusion complete by 8 minutes (Figure 1G,H and Supplementary Figure 1F,G). Interestingly, for SAF-A this biphasic behaviour was not uniform across the cell population. 65.7% of cells showed transient but robust accumulation followed by exclusion, while in the remaining 34.3% the relative fluorescence intensity remained below that of the GFP control indicating exclusion without prior recruitment (Supplementary Figure 1F,G). hnRNPC was the only family member whose distribution was unchanged after microirradiation (Supplementary Figure 1H).

**Figure 1.**
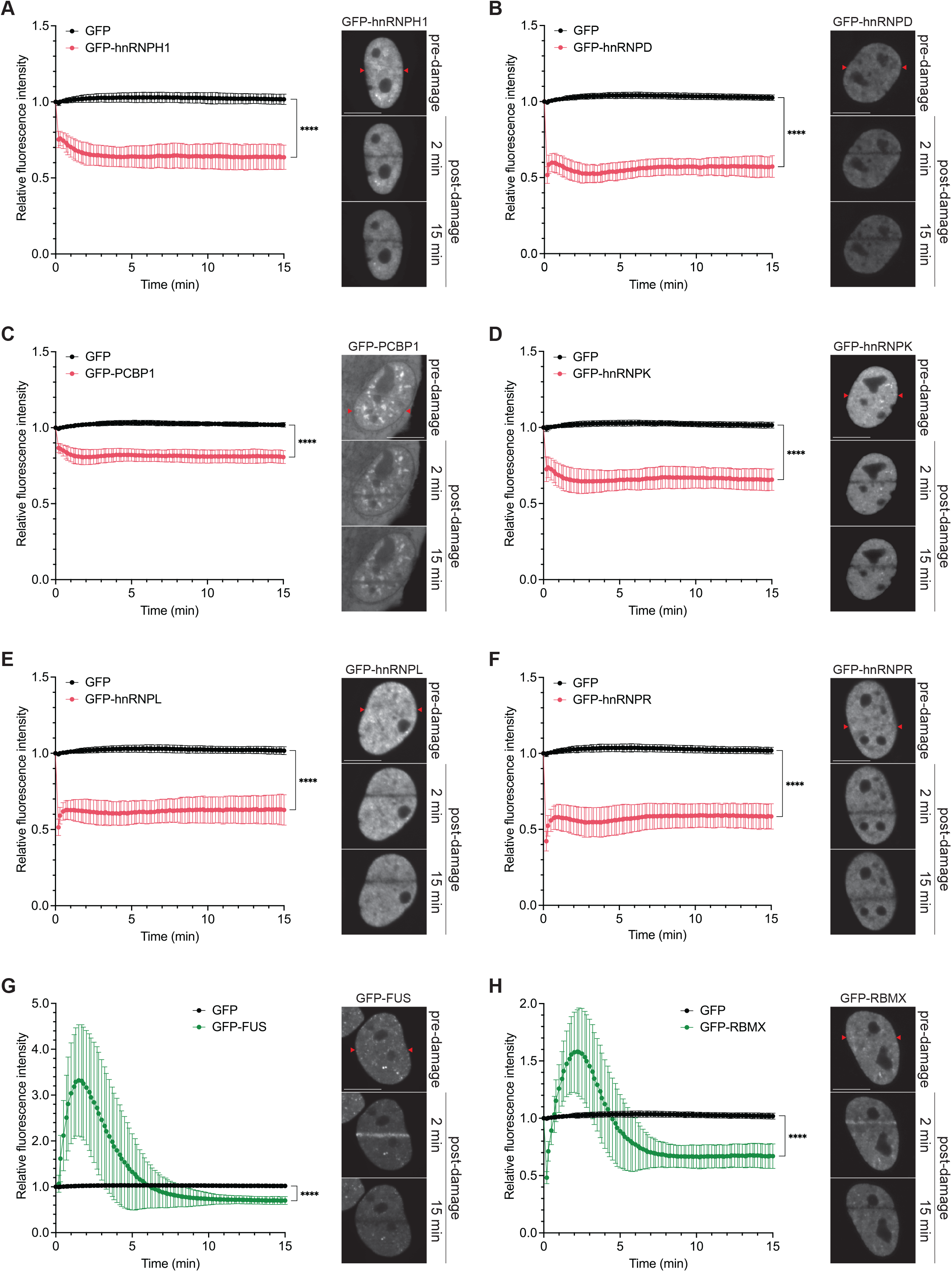
Exclusion from DSBs is a pervasive feature of the hnRNP family. (A-H) HeLa cells transfected with the indicated constructs were subjected to laser microirradiation. Images were obtained at 10-sec intervals for 15 min, and relative fluorescence intensity along the irradiated line was quantified. (A) Dynamics of GFP (n = 38) and GFP-hnRNPH1 (n = 32) at laser-damaged sites (left), and representative images of GFP-hnRNPH1 before and after laser microirradiation (right). (B) Dynamics of GFP (n = 31) and GFP-hnRNPD (n = 31) at laser-damaged sites (left), and representative images of GFP-hnRNPD before and after laser microirradiation (right). (C) Dynamics of GFP (n = 32) and GFP-PCBP1 (n = 31) at laser-damaged sites (left), and representative images of GFP-PCBP1 before and after laser microirradiation (right). (D) Dynamics of GFP (n = 33) and GFP-hnRNPK (n = 36) at laser-damaged sites (left), and representative images of GFP-hnRNPK before and after laser microirradiation (right). (E) Dynamics of GFP (n = 56) and GFP-hnRNPL (n = 34) at laser-damaged sites (left), and representative images of GFP-hnRNPL before and after laser microirradiation (right). (F) Dynamics of GFP (n = 40) and GFP-hnRNPR (n = 37) at laser-damaged sites (left), and representative images of GFP-hnRNPR before and after laser microirradiation (right). (G) Dynamics of GFP (n = 39) and GFP-FUS (n = 33) at laser-damaged sites (left), and representative images of GFP-FUS before and after laser microirradiation (right). (H) Dynamics of GFP (n = 40) and GFP-RBMX (n = 33) at laser-damaged sites (left), and representative images of GFP-RBMX before and after laser microirradiation (right). Data are shown as mean ± SD from 3 independent experiments; two-way ANOVA, ****p < 0.0001. The red arrows indicate the microirradiated area. Scale bars represent 10 µM. Source data are provided as a Source Data file.

Together, these data show that exclusion from damaged chromatin is a general property of the hnRNP family rather than a feature of individual members. This raises the question of whether it is functionally required for DSB signalling or repair.

### hnRNPH1 is persistently excluded from DNA double-strand breaks

hnRNPH1 was among the most strongly excluded family members, and we selected it for further characterisation. It has been linked to DSB repair indirectly through alternative splicing of mouse *Trp53bp1* in meiotic cells (65) and of *PRMT5* in the response to ionising radiation in hepatocellular carcinoma (66), and proteomic studies have identified it in association with RPA (67, 68). Whether hnRNPH1 acts directly at DNA lesions has not been addressed.

We found that hnRNPH1 exclusion was sustained and persists for up to 4 hours after DNA damage induction (Figure 2A,B). Laser microirradiation generates a mix of lesions that include single- and double-strand breaks and base damage, and also damages RNA and proteins along the irradiated track (69). We therefore asked whether hnRNPH1 exclusion is specific to DSBs. To address this question, we used the mCherry-LacI-FOKI U2OS DSB reporter system in which the FOKI endonuclease generates clustered DSBs at a specific genomic locus (44). We observed that GFP-hnRNPH1, but not GFP alone, was excluded from γ-H2AX-marked FOKI-mCherry foci (Figure 2C,D). hnRNPH1 is therefore excluded from site-specific DSBs, independently of the lesion spectrum generated at laser microirradiation tracks.

**Figure 2.**
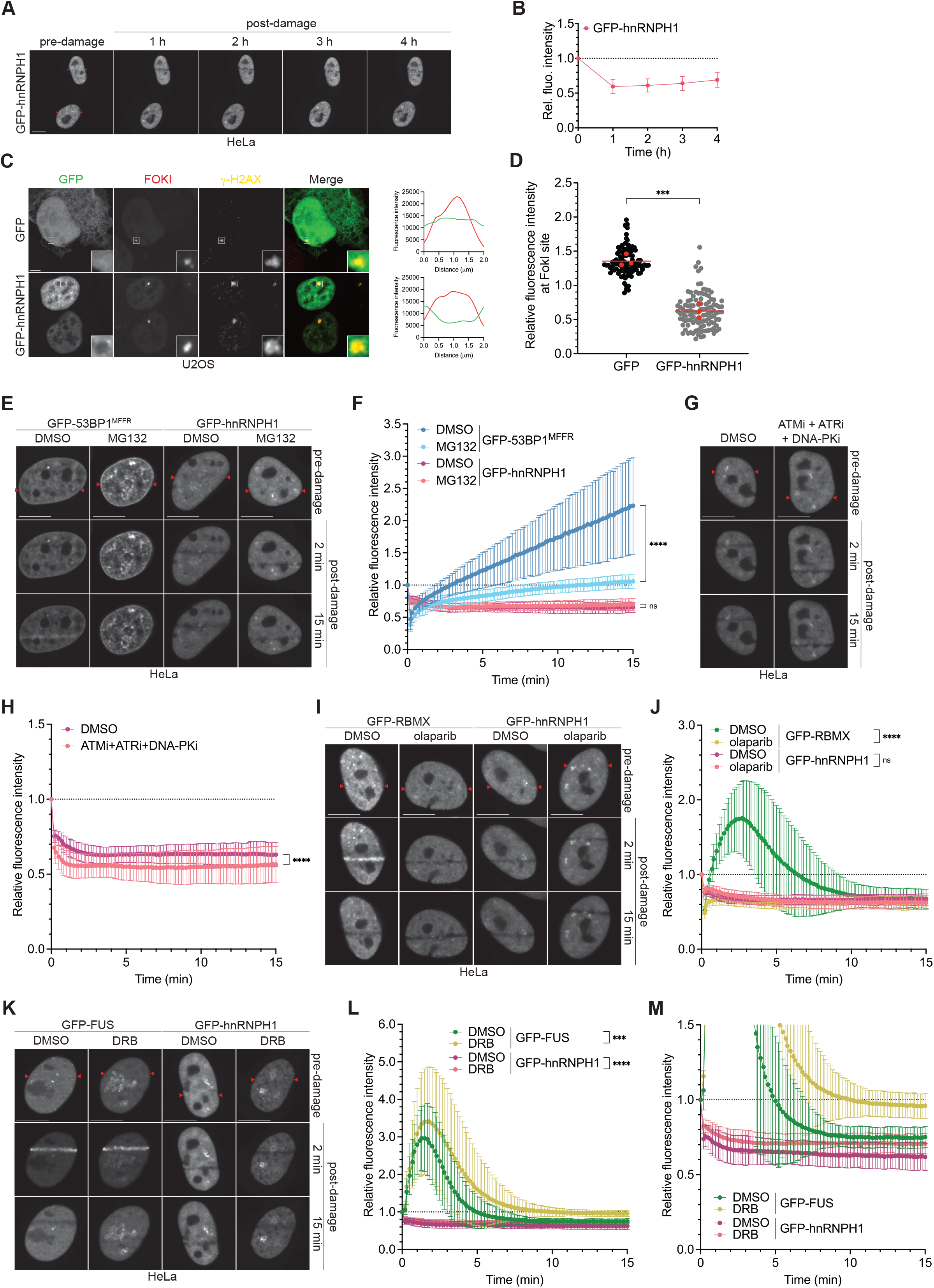
hnRNPH1 is persistently excluded from DNA double-strand breaks. (A) HeLa cells transfected with GFP-hnRNPH1 were laser microirradiated. Images were obtained at 1-hour intervals for 4 hours. The red arrows indicate the microirradiated area. Scale bar represents 5 μm. (B) Quantification of (A). Data are shown as mean ± SD; n = 15. (C) U2OS ER-mCherry-LacI-FOKI-DD cells were transfected with the indicated constructs and DSBs were induced by treatment of these cells with 1 µM Shield-1 and 1 µM 4-OHT for 4 hours. Cells were fixed and processed for γ-H2AX immunofluorescence. Scale bar represents 5 μm. Insets represent 4x magnifications of the indicated fields. A random straight line was drawn across through a single Z section of the nuclei (shown in the Merge images). The fluorescence intensity of GFP and γ-H2AX was quantitated along the length of the line to generate a line profile (right). (D) Quantification of (C). Data are shown as superplots, the bar represents the mean of 3 independent replicates; GFP (n = 102), GFP-hnRNPH1 (n = 113); unpaired t-test, ***p < 0.001. (E) HeLa cells were transfected with GFP-53BP1^MFFR^ or GFP-hnRNPH1, and were treated with DMSO or 25 μM MG132 for 1.5 hours prior to laser microirradiation. The red arrows indicate the microirradiated area. Scale bars represent 10 µM. (F) Quantification of (E). Data are shown as mean ± SD from 3 independent experiments; GFP-53BP1^MFFR^+DMSO (n = 30), GFP-53BP1^MFFR^+MG132 (n = 41), GFP-hnRNPH1+DMSO (n = 30), GFP-hnRNPH1+MG132 (n = 36); two-way ANOVA, n.s. non-significant (p ≥ 0.05), ****p < 0.0001. (G) HeLa cells were transfected with GFP-hnRNPH1, and were treated with a combination of 20 μM ATM inhibitor (KU-55933), 10 μM ATR inhibitor (VE-821) and 10 μM DNA-PK inhibitor (NU-7441) for 1 hour prior to laser microirradiation. The red arrows indicate the microirradiated area. Scale bars represent 10 µM. (H) Quantification of (G). Data are shown as mean ± SD from 3 independent experiments; GFP-hnRNPH1+DMSO (n = 35), GFP-hnRNPH1+ATMi+ATRi+DNA-PKi (n = 44); two-way ANOVA, ****p < 0.0001. (I) HeLa cells were transfected with GFP-RBMX or GFP-hnRNPH1, and were treated with 10 μM PARP inhibitor (olaparib) for 1 hour prior to laser microirradiation. The red arrows indicate the microirradiated area. Scale bars represent 10 µM. (J) Quantification of (I). Data are shown as mean ± SD from 3 independent experiments; GFP-RBMX+DMSO (n=36), GFP-RBMX+olaparib (n=34), GFP-hnRNPH1+DMSO (n = 39), GFP-hnRNPH1+olaparib (n = 34); two-way ANOVA, n.s. non-significant (p ≥ 0.05), ****p < 0.0001. (K) HeLa cells were transfected with GFP-FUS or GFP-hnRNPH1, and were treated with 100 μM transcription inhibitor (DRB) for 3 hours prior to laser microirradiation and during live-cell imaging. The red arrows indicate the microirradiated area. Scale bars represent 10 µM. (L) Quantification of (K). Data are shown as mean ± SD from 3 independent experiments; GFP-FUS+DMSO (n = 30), GFP-FUS+DRB (n = 32), GFP-hnRNPH1+DMSO (n = 36), GFP-hnRNPH1+DRB (n = 39); two-way ANOVA, ***p < 0.001, ****p < 0.0001. (M) Zoomed-in view of (L). Source data are provided as a Source Data file.

We next asked which signal triggers hnRNPH1 exclusion. Importantly, exclusion did not reflect protein degradation, as we detected no change in total hnRNPH1 levels following ionising radiation (IR) (Supplementary Figure 2A). We then reasoned that exclusion might instead be driven by the post-translational modifications that propagate DNA damage signalling, ubiquitylation, phosphorylation and poly(ADP)-ribosylation. Exclusion persisted when we inhibited ubiquitin-dependent signalling at breaks with the proteasome inhibitor MG132 (25 μM for 1.5 hours), although 53BP1 recruitment was impaired under the same conditions as expected (Figure 2E,F) (14). Similarly, we saw no effect when we inhibited ATM, ATR and DNA-PK simultaneously for 1 hour prior to laser microirradiation (Figure 2G,H, Supplementary Figure 2B). Because DYRK1A is another kinase that has been linked to the chromatin-based DSB response, we also tested its involvement (70–72). However, hnRNPH1 exclusion was unaffected by treatment with DYRK1A inhibitors Harmine (10 μM) or INDY (25 μM) for 24 hours before DNA damage induction (Supplementary Figure 2C,D). Finally, PARP1 activity is required for the recruitment of several hnRNPs to DSBs, including RBMX, CIRBP, FUS and SAF-A (51, 53, 54, 58). We therefore used RBMX as a positive control for PARP inhibition. Notably, while olaparib treatment (10 μM, 1 hour prior to laser microirradiation) effectively abolished RBMX recruitment to laser microirradiation tracks, it left the subsequent exclusion of RBMX intact and also had no effect on hnRNPH1 exclusion (Figure 2I,J). hnRNP recruitment and exclusion are therefore separable events, and exclusion does not depend on PAR synthesis.

Exclusion need not be driven by damage signalling. For SAF-A and FUS, it has instead been attributed to transcriptional shutdown at the break site (51, 58). This prompted us to ask whether the same applies to hnRNPH1 exclusion. Inhibition of transcription with the CDK9 inhibitor DRB (100 μM, 3 hours prior to laser microirradiation as well as during live-cell imaging) indeed prevented FUS exclusion from laser microirradiation tracks, but had no effect on hnRNPH1 (Figure 2K-M and Supplementary Figure E). hnRNPH1 exclusion is thus not a consequence of local, DSB-induced transcriptional inhibition (44).

Together, these results show that hnRNPH1 is excluded from DSBs independently from the lesion spectrum generated by microirradiation and that exclusion is sustained over several hours. Exclusion is not driven by protein turnover, DNA damage signalling or transcriptional shutdown, suggesting that recruitment to and exclusion from can be driven by distinct mechanisms.

### The GYR domain of hnRNPH1 which promotes LLPS is important for exclusion

Having found no evidence that hnRNPH1 exclusion is triggered by DNA damage signalling, we asked whether exclusion is driven by features of hnRNPH1 itself. hnRNPH1 is a 449-amino acid protein containing three quasi-RNA recognition motifs (qRRMs) that mediate RNA binding. In addition, it contains two low complexity domains, a glycine-tyrosine-arginine-rich region (GYR domain) and a C-terminal glycine-rich region (GY domain) (Figure 3A). The GYR domain mediates liquid–liquid phase separation (LLPS) and the formation of hydrogel droplets *in vitro* (73). We generated GFP-tagged deletion mutants lacking each domain individually or lacking all three qRRMs (Figure 3A and Supplementary Figure 3A). All mutants were predominantly nuclear. GFP-hnRNPH1-ΔGYR showed some additional signal, consistent with the removal of the Nuclear Localisation Signal (NLS), but the majority of the protein remained nuclear (Figure 3D).

**Figure 3.**
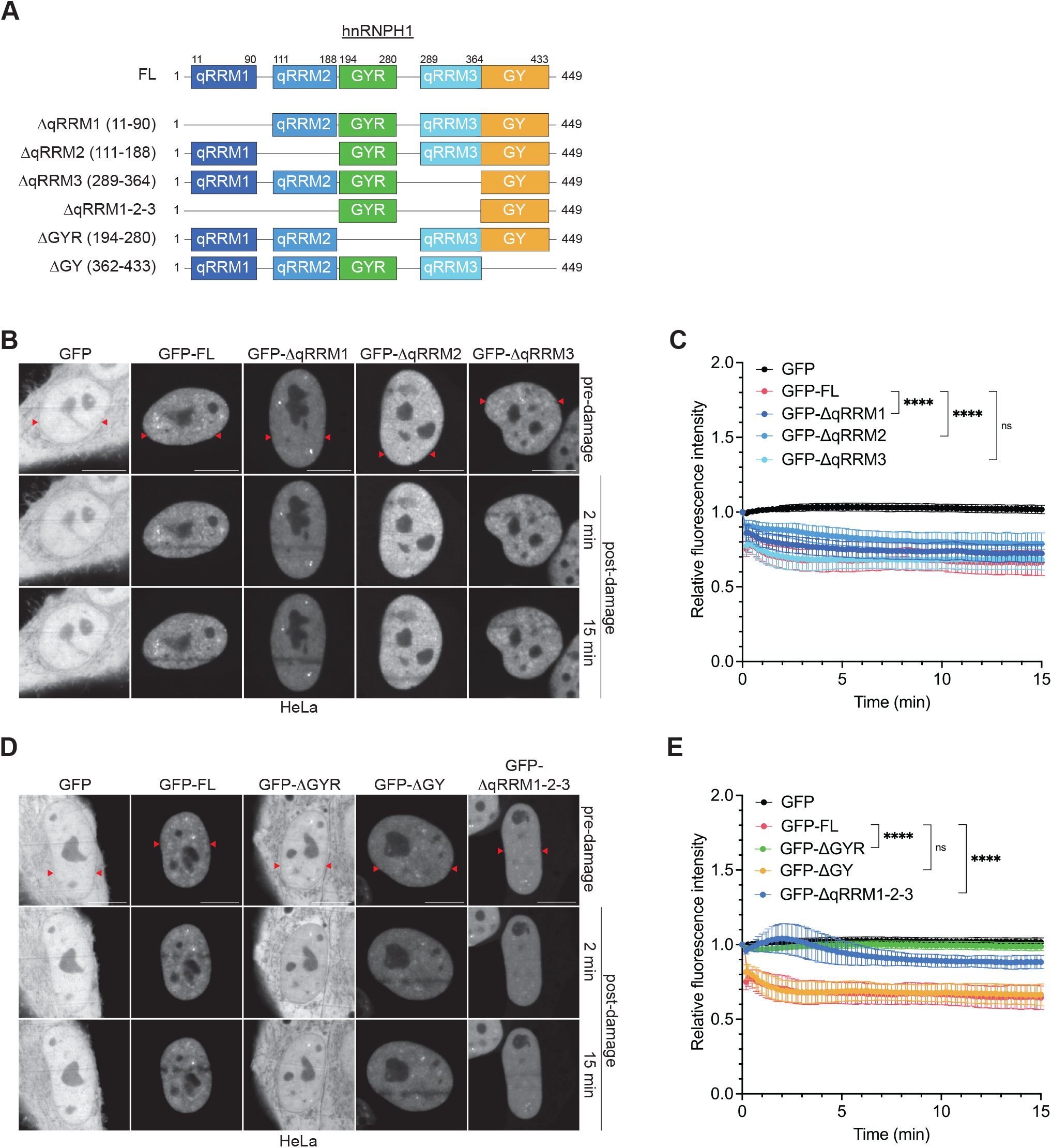
The GYR domain of hnRNPH1 which promotes LLPS is important for exclusion. (A) hnRNPH1 domains architecture and variants used. hnRNPH1 contains three quasi-RNA Recognition Motifs (qRRMs), a Glycine-Tyrosine-Arginine rich domain (GYR) and a Glycine-Tyrosine rich domain (GY). FL: full-length. (B) HeLa cells were transfected with the indicated constructs and were subjected to laser microirradiation. The red arrows indicate the microirradiated area. Scale bars represent 10 µM. (C) Quantification of (B). Data are shown as mean ± SD from 3 independent experiments; GFP (n = 30), GFP-FL (n = 30), GFP-ΔqRRM1 (n = 31), GFP-ΔqRRM2 (n = 31) and GFP-ΔqRRM3 (n = 34); two-way ANOVA, n.s. non-significant (p ≥ 0.05), ****p < 0.0001. (D) HeLa cells were transfected with the indicated constructs and were subjected to laser microirradiation. The red arrows indicate the microirradiated area. Scale bars represent 10 µM. (E) Quantification of (D). Data are shown as mean ± SD from 3 independent experiments; GFP (n = 35), GFP-FL (n = 34), GFP-ΔGYR (n = 36), GFP-ΔGY (n = 32), and GFP-ΔqRRM1-2-3 (n = 35); two-way ANOVA, n.s. non-significant (p ≥ 0.05), ****p < 0.0001. Source data are provided as a Source Data file.

We found that the deletion of any single qRRM did not impair hnRNPH1 exclusion from DNA damage (Figure 3B,C), indicating that no individual RNA-binding domain is required. Notably, the GFP-hnRNPH1-ΔqRRM1-2-3 mutant behaved differently. It showed a weak wave of hnRNPH1 recruitment followed by delayed exclusion that was weaker than for the single GFP-hnRNPH1-ΔqRRM mutants (Figure 3D,E). Removal of all three modules together therefore alters the dynamics of hnRNPH1 at DSBs, although exclusion still occurs.

Among the two low-complexity domains, GFP-hnRNPH1-ΔGY was excluded to the same extent as full-length hnRNPH1 (Figure 3D,E). In contrast, GFP-hnRNPH1-ΔGYR was not excluded from laser microirradiation tracks and showed the weakest response of all mutants tested (Figure 3D,E). The GYR domain is therefore required for hnRNPH1 exclusion from DSBs.

Together, these results suggest that hnRNPH1 exclusion depends on the GYR low-complexity domain, which mediates phase separation (73), with RNA binding likely contributing redundantly through the three qRRMs.

### Tethering hnRNPH1 to DSBs impairs 53BP1 recruitment

To test whether exclusion is functionally required for the DSB response, we tethered hnRNPH1 to damaged chromatin by fusing it to the forkhead-associated (FHA) domain of RNF8, which directs proteins to phosphorylated MDC1 at DSBs (Figure 4A and Supplementary Figure 4A) (13, 14, 16, 74). This strategy has been used in multiple studies to target proteins to damaged chromatin (27, 75, 76). As a control we used the phospho-binding-deficient FHA^R42A^ mutant, which does not localise to DNA breaks (13, 14). GFP-FHA and GFP-FHA-hnRNPH1 both localised to IR-induced foci 1 hour post-IR while GFP-FHA^R42A^-hnRNPH1 failed to localise (Figure 4B).

**Figure 4.**
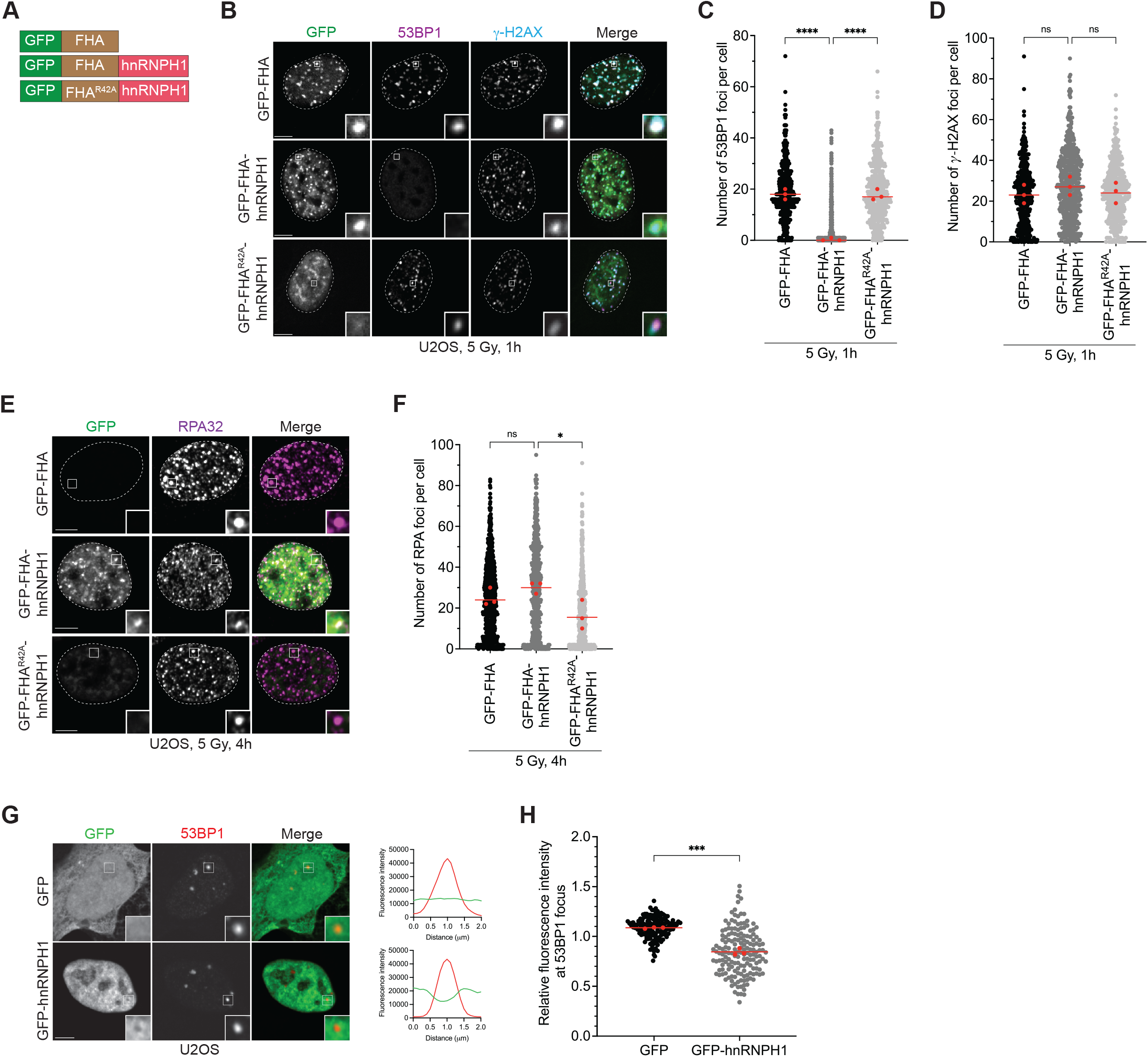
Tethering hnRNPH1 to DSBs impairs 53BP1 recruitment. (A) GFP-FHA, GFP-FHA-hnRNPH1 and GFP-FHA^R42A^-hnRNPH1 fusion proteins. (B) U2OS cells stably expressing the indicated constructs were irradiated (5 Gy), fixed 1 hour after irradiation and processed for 53BP1 and γ-H2AX immunofluorescence. Scale bar represents 5 μM. Dashed lines indicate nucleus outlines (as determined using DAPI staining; not shown). Insets represent 4x magnifications of the indicated fields. (C) Quantification of 53BP1 foci per cell (from B). Data are shown as superplots, the bar represents the median of 3 independent replicates; at least 100 cells per condition were counted; one-way ANOVA, ****p < 0.0001. (D) Quantification of γ-H2AX foci per cell (from B). Data are shown as superplots, the bar represents the median of 3 independent replicates; at least 100 cells per condition were counted; one-way ANOVA, n.s. non-significant (p ≥ 0.05). (E) U2OS cells stably expressing the indicated constructs were irradiated (5 Gy), fixed 4 hours after irradiation and processed for RPA32 immunofluorescence. Scale bar represents 5 μM. Dashed lines indicate nucleus outlines (as determined using DAPI staining; not shown). Insets represent 4,3x magnifications of the indicated fields. (F) Quantification of (E). Data are shown as superplots, the bar represents the median of 3 independent replicates; at least 100 cells per condition were counted; one-way ANOVA. n.s. non-significant (p ≥ 0.05), *p < 0.05. (G) U2OS cells were transfected with the indicated constructs and processed for 53BP1 immunofluorescence. Scale bar represents 5 μM. Insets represent 4x magnifications of the indicated fields. A random straight line was drawn across through a single Z section of the nuclei (shown in the Merge images). The intensity of GFP and 53BP1 was quantitated along the length of the line to generate a line profile (right). (H) Quantification of (G). Data are shown as superplots, the bar represents the mean of 3 independent replicates; GFP (n = 149), GFP-hnRNPH1 (n = 165); unpaired t-test, ***p < 0.001. Source data are provided as a Source Data file.

We found that, compared with GFP-FHA^R42A^-hnRNPH1, recruitment of GFP-FHA-hnRNPH1 strongly suppressed 53BP1 focus formation after IR (Figure 4B,C, Supplementary Figure 5A). This effect was specific to 53BP1 as both γ-H2AX and MDC1 foci formed normally (Figure 4B,D, Supplementary Figure 4B,C and Supplementary Figure 5B,C), indicating that upstream signalling at DSBs was intact. Notably, RPA focus formation was equally unaffected, suggesting that DNA end-resection proceeds normally in the presence of chromatin-tethered hnRNPH1 (Figure 4E,F and Supplementary Figure 5D). Importantly, total 53BP1 levels were unchanged in cells expressing the FHA constructs, excluding that the phenotype is caused by changes in 53BP1 protein levels (Supplementary Figure 4D).

Finally, we asked whether the relationship between hnRNPH1 and 53BP1 is also apparent without exogenous DNA damage. Indeed, in unchallenged cells, GFP-hnRNPH1 was excluded from 53BP1 bodies (Figure 4G,H) (77, 78), indicating that hnRNPH1 exclusion and 53BP1 accumulation are coupled under physiological conditions, and that the phenotype observed by chromatin tethering is not an artefact of forced local retention.

Together, these results demonstrate that retention of hnRNPH1 at damaged chromatin selectively interferes with 53BP1 accumulation. Upstream signalling and DNA end-resection are unaffected, suggesting that hnRNPH1 does not act by non-specifically disrupting focus assembly, but at the step at which 53BP1 engages damaged chromatin.

## DISCUSSION

The systematic comparison reported here establishes exclusion as a pervasive feature of hnRNP family members at DSBs. While exclusion has been reported for individual hnRNPs, its scope across the family had not been examined (51–59). We find that 13 of 14 core members we tested were cleared from damaged chromatin, 10 directly and 3 following a brief phase of recruitment. The recruitment described for individual hnRNPs is therefore neither typical of the family nor sustained (51–56, 58, 59). Exclusion occurs within minutes, rapid enough to coincide with the earliest steps of the DSB response. In the case of hnRNPH1, it is sustained for several hours and has direct functional consequences, since forcing hnRNPH1 to remain at breaks selectively impairs 53BP1 accumulation. Removal of hnRNPs is therefore an early and consequential part of the chromatin-based DSB response.

hnRNPH1 tethering affected neither γ-H2AX and MDC1 accumulation nor RPA loading. The DSB is therefore recognised, the signalling marks set by the DSB response are deposited, and only 53BP1 fails to accumulate. Notably, these factors differ in how they engage damaged chromatin. MDC1 binds γ-H2AX through a defined BRCT interaction and RPA binds end-resected ssDNA directly (15, 79), whereas 53BP1 accumulates through weak, multivalent engagement of a bivalent histone mark that is further stabilised through oligomerisation (28). Assembly of this kind depends on the local concentration of competing multivalent proteins, while higher-affinity, stoichiometric binding does not (80). Consistent with this, 53BP1 assembly at breaks is sensitive to perturbations of the nuclear environment that leave γ-H2AX and MDC1 accumulation intact (35). hnRNPs are abundant and highly multivalent and, in the context of damaged chromatin, might interfere with the weak interactions through which 53BP1 accumulates (50). Notably, the GYR domain is both the region that is required for hnRNPH1 exclusion and the region that drives its phase separation *in vitro* (73). We therefore propose that hnRNPs must be cleared from damaged chromatin for the assembly of repair factors that accumulate by high avidity, rather than repair factor assembly in general.

Exclusion can likely be achieved through multiple mechanisms. For FUS, exclusion has recently been shown to require the transcription-associated kinases CDK7, CDK9 and CDK12/13 as well as the RNA polymerase II subunit RPB1, which is itself excluded from lesions with similar kinetics (58). This is consistent with earlier work attributing SAF-A and FUS exclusion to transcriptional shutdown at DSB sites (51). We found no evidence that hnRNPH1 depends on the transcription-associated mechanism described for FUS, and the signal that drives its exclusion remains unidentified. However, hnRNPH1 exclusion does require the GYR low-complexity domain and, to a lesser degree, the qRRMs, which potentially points away from a specific signalling trigger and towards a property of the protein itself. In addition, exclusion is also mechanistically separable from recruitment. In the case of RBMX, olaparib abolished recruitment to damaged chromatin but left the subsequent exclusion intact, showing within a single protein that the two behaviours are independently controlled.

Exclusion and local transcriptional repression are not ordered the same way for all family members. Exclusion of FUS follows transcriptional shutdown (51, 58), whereas release of hnRNPA2B1 from damaged chromatin is itself what destabilises RNA polymerase II and represses local transcription (61). hnRNPH1 exclusion was unaffected by DRB, which likely places it outside the transcription-coupled route described for FUS. Removing all three qRRMs did, however, affect exclusion, suggesting that it depends on RNA binding but not on RNA polymerase II elongation at the break site. Because deleting the qRRMs of hnRNPH1 removes RNA engagement throughout the nucleus rather than only at damaged chromatin, we hypothesise that RNA binding determines the distribution and residence time of hnRNPH1 in the nucleoplasm rather than retention at DSBs.

Two, not mutually exclusive mechanisms could explain the specificity of the 53BP1 phenotype. The GYR domain required for hnRNPH1 exclusion has been shown to drive liquid-liquid phase separation *in vitro* (73), and 53BP1 itself assembles into compartments with liquid-like behaviour at breaks (35). If 53BP1 accumulation depends on the material properties of damaged chromatin in a way that MDC1 binding and RPA loading do not, a retained low-complexity protein such as hnRNPH1 could be expected to interfere with 53BP1 selectively. Alternatively, the effect may be mediated by a defined interaction between hnRNPH1, the long non-coding RNA NEAT1 and the NEAT1-associated factor TIRR. TIRR occludes the 53BP1 Tudor domain and blocks its binding to H4K20me2, and NEAT1 relieves this inhibition by competing with 53BP1 for TIRR (81). Because hnRNPH1 binds NEAT1 (82), its retention at damaged chromatin could reduce the pool of NEAT1 molecules available to TIRR, leaving 53BP1 inhibited (83).

More broadly, our results suggest that retaining hnRNPH1 at damaged chromatin is disruptive, which is not the same as showing that removal is required. Distinguishing the two concepts is the central question raised by these data, and it applies to the hnRNP family as a whole: whether exclusion contributes to DNA repair or is a consequence of it. What our findings establish is that a DSB arises in a nucleus that is densely populated by RNA-binding proteins such as the hnRNPs, and that the response to it involves clearing them from the damage site as well as recruiting repair factors. Thus, the composition of the chromatin surrounding a break site is defined as much by what has been cleared from it as by what has arrived.

## MATERIAL AND METHODS

### Cell culture

HeLa 1.2.11, U2OS FlpIn and U2OS ER-mCherry-LacI-FOKI-DD cells were cultured in Dulbecco’s Modified Eagle Medium (DMEM) with Glutamax (Life Technologies) supplemented with 10% (v/v) fetal bovine serum (FBS, Life Technologies). The inducible GFP-FHA, GFP-FHA-hnRNPH1 and GFP-FHA^R42A^-hnRNPH1 cell lines were generated using the Flp-In T-REx system (Invitrogen) as described in the manufacturer’s protocol. Each construct was cloned into the pcDNA5-FRT/TO-GFP vector followed by co-transfection with the pOG44 vector (Flp recombinase) into U2OS FlpIn host cell lines. The host cell line was cultured in DMEM supplemented with 15,5 μg/ml zeocin (Invitrogen). Recombination events were selected with 250 μg/ml Hygromycin B (ThermoScientific). U2OS FlpIn stable cell lines were cultured in DMEM supplemented with Blasticidin S (5 mg/ml, Life Technologies) and Hygromycin B (200 mg/ml, Sigma-Aldrich). Cells were maintained using standard tissue culture procedures and kept in a humidified incubator set to 37°C and 5% CO_2_. Cells were routinely tested for mycoplasma contamination. All plasmids were transfected using Effectene Transfection Reagent (Qiagen, 301425) according to the manufacturer’s protocol.

### Cloning

All DNA preparations (including agarose gel extractions, minipreps and maxipreps) were done with DNA purification kits from QIAGEN according to the manufacturer’s instructions. All constructs were confirmed by sequencing.

The cDNA of hnRNPH1, PTBP1, FUS, RBMX and hnRNPR was amplified by PCR from an IMR-32 cDNA library, and hnRNPQ from a HeLa cDNA library. The cDNA of hnRNPD, hnRNPE1, hnRNPK and hnRNPU were amplified from a HEK293 cDNA library. The cDNA of hnRNPL was synthesised with the GeneArt service from Thermofisher. The cDNA of hnRNPF was amplified from the p3x-Flag-hnRNPF plasmid (Addgene #21926). The cDNA of hnRNPA2B1 was amplified by PCR from the MBP-hnRNPA2_FL_WT plasmid (Addgene #98662). The cDNAs of hnRNPC1 and hnRNPC2 were amplified from the pcDNA3.1-hnRNPC-myc plasmid (Addgene #71661) and HeLa cDNA library, respectively. The cDNA of the 53BP1-MFFR region was a kind gift of Amélie Fradet-Turcotte (Canada). All cDNAs were ligated into pcDNA5-FRT/TO-GFP.

To generate hnRNPH1 fragments and mutants, standard protocols for primer-directed mutagenesis or self-ligation of truncated PCR-products were used. The details of the internal pcDNA5-FRT/TO-GFP-hnRNPH1 deletions are as follows: GFP-ΔqRRM1, amino acid residues 11-90; GFP-ΔqRRM2, amino acid residues 111-188; GFP-ΔqRRM3, amino acid residues 289-364; GFP-ΔGYR, amino acid residues 194-280; GFP-ΔGY, amino acid residues 362-433; GFP-ΔqRRM1-2-3, amino acid residues 11-90, 111-188 and 289-364.

To generate RNF8-FHA fusions, the N-terminus of RNF8 (amino acid residues 1-160) was synthesised with the DNA synthesis service from Twist Bioscience, PCR amplified and inserted into pcDNA5-FRT/TO-GFP and pcDNA5-FRT/TO-GFP-hnRNPH1. To generate FHA^R42A^ mutant, standard protocols for primer-directed mutagenesis was used.

### Laser microirradiation

75,000 cells were seeded on μ-dish 35 mm high glass bottom dishes (Ibidi #81158). The day after, cells were transfected with 200-300 ng of pcDNA5-FRT/TO-GFP plasmids, using Effectene Transfection Reagent (Qiagen, 301425) and 10 μM 5-bromo-2’-deoxyuridine (BrdU, Sigma-Aldrich). 6 hours post-transfection, media was replaced with fresh medium containing 10 μM BrdU. The day after, immediately prior to imaging, growth medium was replaced with CO_2_-independent Live Cell Imaging medium (Invitrogen #A1429DJ) supplemented with 10 % FBS, 2 mM L-glutamine (100X GlutaMAX).

Laser microirradiation was performed with a spinning disc confocal Andor Dragonfly inverted microscope equipped with a 37°C incubation chamber and a 405 nm laser diode focused through a 60X oil objective. Laser settings (20 μm stripe, 7 mW output, 10 iterations, accurate mode) were chosen to generate a detectable DDR restricted to the laser path in a pre-sensitization-dependent manner without major cytotoxic effects. Laser power was measured each time to reach 7 mW. Images were acquired every 10 or 30 seconds for 15 min.

Analysis of the image sequences was performed by manually segmenting the irradiated area as well as the whole nucleus on Fiji. Mean fluorescence intensities with the irradiated area was background subtracted, divided to the mean nuclear intensity to correct for imaging photobleaching, and then normalised to the signal prior to DNA damage.

### DNA tethering

Cells were grown on #1.5 glass coverslips in a 24-well plate. The expression of GFP-FHA, GFP-FHA-hnRNPH1 and GFP-FHA^R42A^-hnRNPH1 was induced with Doxycyclin (Sigma-Aldrich). 24 hours after Doxycyclin treatment, cells were irradiated (5 Gy) with a Biobeam 8000 and fixed at different time-points after irradiation. Cells were washed with 1X PBS and pre-extracted with ice-cold CSK buffer (20 mM HEPES pH 7.5, 100 mM NaCl, 3 mM MgCl2, 300 mM sucrose, 0.5 % Triton X-100, phosphatase and protease inhibitor tablets) for 2 min on ice. The pre-extraction buffer was removed and cells were immediately fixed with 4 % (v/v) formaldehyde (Life Technologies) in CSK buffer for 30 min at RT.

### FOKI-mediated DNA damage induction

U2OS ER-mCherry-LacI-FOKI-DD cells were seeded on 24-well plates containing glass coverslips without any induction of FOKI. 24 hours after seeding, cells were transfected using 200 ng of GFP or GFP-hnRNPH1 expression vectors. The day after, cells were treated for 4 hours with 1 µM of Shield-1 and 1 µM of 4-Hydroxytamoxifen to promote FOKI stabilization and nuclear translocation. Cells were fixed with 2 % formaldehyde for 20 min at RT and processed as described below.

All intensity analysis was performed on unprocessed images using Fiji. For each cell, an mCherry-FOKI focus was used to generate a mask. The relative GFP fluorescence intensity in the mCherry-FOKI focus mask was calculated by the following formula: RFI^GFP^ = (I_mCherry-FOKI mask_ - I_b_) / (I_nucleus_-I_b_), where I_mCherry-FOKI mask_ is the mean GFP fluorescence intensity at the mCherry-FOKI focus mask, I_b_ is the mean fluorescence intensity of the background, and I_nucleus_ is the mean fluorescence intensity of the nucleus.

### Immunofluorescence

Cells were washed with 1X PBS and pre-extracted with ice-cold pre-extraction buffer (20 mM HEPES pH 7.5, 20 mM NaCl, 5 mM MgCl2, 300 mM sucrose, 0.5 % NP-40, 1 mM DTT, phosphatase and protease inhibitor tablets) for 20 min on ice. The pre-extraction buffer was removed and cells were immediately fixed with 2 % (w/v) formaldehyde (Life Technologies) in PBS for 20 min at RT. After fixation, cells were washed with 1X PBS three times for 5 min respectively and then blocked with Antibody Dilution Buffer (ADB; 10 % (v/v) normal goat serum, 0.1 % (v/v) Triton X-100, 0.1 % (v/v) saponin in PBS) for 1 hour. Cells were incubated with the appropriate primary antibody (see list of antibodies) diluted in ADB overnight at 4°C. After incubation in primary antibodies, cells were washed once with 1X PBS + 0.1 % (v/v) Triton X-100 for 5 min, followed by two washes with 1X PBS for 5 min. Each coverslip was then counterstained with Alexa Fluor secondary antibodies (Thermo Fisher Scientific) diluted 1:1000 in ADB (see list of antibodies), for 1 hour at RT in the dark. After washing once with 1X PBS + 0.1 % (v/v) Triton X-100 for 5 min, cells were washed with 1X PBS two times for 5 min. To visualise nuclei, cells were stained with 0.1 μg/mL DAPI in 1X PBS for 5 min, followed by two washes with 1X PBS. Coverslips were mounted onto glass slides with Prolong Diamond Antifade mounting agent (Life Technologies) and left to dry overnight at RT in the dark.

For the quantification of GFP and GFP-hnRNPH1 relative fluorescence intensity at 53BP1 foci, each 53BP1 focus was used to generate a mask. The relative GFP fluorescence intensity in the 53BP1 focus mask was calculated by the following formula: RFI^GFP^ = (I_53BP1 mask_ - I_b_) / (I_nucleus_-I_b_), where I_53BP1 mask_ is the mean GFP fluorescence intensity at the 53BP1 focus mask, I_b_ is the mean fluorescence intensity of the background, and I_nucleus_ is the mean fluorescence intensity of the nucleus.

### Image acquisition and analysis of IF data

Images of immunofluorescence staining were acquired with a spinning disc confocal Andor Dragonfly inverted microscope equipped with a 60X oil objective. Following acquisition, images were imported into Fiji for quantification. For the tethering experiments, the analysis was performed only on GFP-positive cells detected by the Ilastik software, and automated quantification of foci was performed with a custom script in Fiji.

### Western blotting

Cells were lysed using cell lysis buffer (50 mM Hepes-KOH, pH 7.5, 100 mM KCl, 2 mM EDTA, 0.5 % (v/v) IGEPAL, 10 % (v/v) glycerol, 1 mM DTT, 1× protease and phosphatase inhibitors). Lysates were cleared through centrifugation at 13,000× *g* for 20 min at 4 °C. Protein concentration was determined using the BCA method (DC protein assay (Biorad)) according to the manufacturer’s instructions. Lysates were denatured in 1× Laemmli Sample Buffer (Biorad, 1610747) and 5 mM DTT for 5 min at 95°C. Alternatively, cells were lysed using 1x Laemmli Sample Buffer supplemented with 1 mM DTT and Benzonase for 1 hour on ice, and boiled for 5 min at 95°C. Equal amount of proteins were subjected to SDS-PAGE on Tris-Glycine gel. Proteins were transferred onto a nitrocellulose membrane (Biorad) at 100 V for 2 hours. Membranes were blocked either in 5% non-fat dried milk or 5% BSA in TBS containing 0.1% Tween-20 buffer for at least 1 hour at RT and subsequently incubated overnight at 4 °C with primary antibody (see list of antibodies). Vinculin, Tubulin or GAPDH was used as a loading control. HRP-conjugated anti-mouse secondary antibody or HRP-conjugated anti-rabbit secondary antibody were used. Immune complexes were detected with ECL Western Blotting Reagent (Thermo Fisher Scientific). Blots were analysed using Image Lab v6.0 (Biorad).

### Quantification and statistical analysis

All data analysis was performed using Microsoft Excel, GraphPad Prism and/or Fiji. Differences were analysed with several statistical tests, as indicated in figure legends. In all graphs, the mean or median ± SD (standard deviation) is plotted, as indicated in figure legends. Significance; ns, not significant (p ≥ 0.05), *p < 0.05, **p < 0.01, ***p < 0.001, ****p < 0.0001.

### Reagents

| **Antibody target** | **Species** | **Supplier** | **RRID** | **Application** |
|---|---|---|---|---|
| 53BP1 | rabbit | Biomol, A300-272A | AB_185520 | IF (1:1000) |
| γ-H2AX | mouse | Sigma-Aldrich, ZMS05636 | AB_2924829 | IF (1:10,000) |
| γ-H2AX | rabbit | Cell Signalling Technology, 2577S | AB_2118010 | IF (1:1000), WB (1:5000) |
| H2AX | rabbit | Abcam, ab11175 | AB_297814 | WB (1:5000) |
| MDC1 | rabbit | Abcam, ab11171 | AB_297810 | IF (1:500) |
| RPA32 | mouse | Abcam, ab2175 | AB_302873 | IF (1:500) |
| Anti-mouse AF647 | mouse | Invitrogen, A-21236 | AB_2535805 | IF (1:1000) |
| Anti-mouse AF546 | mouse | Invitrogen, A-11030 | AB_144695 | IF (1:1000) |
| Anti-rabbit AF647 | rabbit | Invitrogen, A-21245 | AB_2535813 | IF (1:1000) |
| Anti-rabbit AF546 | rabbit | Invitrogen, A-11035 | AB_143051 | IF (1:1000) |
| hnRNPH1 | rabbit | ProteinTech, 14774-1-AP | AB_2120339 | WB (1:1000) |
| pCHK2 | rabbit | Cell Signalling Technology, 2197 | AB_2080501 | WB (1:1000) |
| CHK2 | mouse | Merck Millipore, 05-649 | AB_2244941 | WB (1:1000) |
| GFP | mouse | Sigma, 11814460001 | AB_390913 | WB (1:1000) |
| RPB1-pSer5-CTD | rat | Chromotek, 3E8 | AB_2631404 | WB (1:500) |
| RPB1-pSer2-CTD | rat | Chromotek, 3E10 | AB_2631403 | WB (1:500) |
| RPB1-pSer7-CTD | rat | Chromotek, 4E12 | AB_2631443 | WB (1:500) |
| RPB1-NTD | rabbit | Cell Signalling Technology, 14958S | AB_2687876 | WB (1:1000) |
| Vinculin | mouse | Sigma, V4505 | AB_477617 | WB (1:1000) |
| Tubulin | mouse | Sigma, T6074 | AB_477582 | WB (1:1000) |
| GAPDH | rabbit | Cell Signalling Technology, 2118S | AB_561053 | WB (1:1000) |
| HRP-conjugated anti-mouse secondary antibody | horse | Cell Signalling Technology, 7076 | AB_330924 | WB (1:5000) |
| HRP-conjugated anti-rabbit secondary antibody | goat | Cell Signalling Technology, 7074 | AB_2099233 | WB (1:5000) |
| **Chemical** | **Purpose** | **Supplier** | **Condition of use<sup>a</sup>** |  |
| MG132 | Proteasome inhibitor | Sigma-Aldrich, M7449 | 25 µM for 1.5 h |  |
| Olaparib | PARP inhibitor | Biomol, Cay10621 | 10 µM for 1 h |  |
| KU-55933 | ATM inhibitor | Sigma-Aldrich, SML1109 | 20 µM for 1 h |  |
| VE-821 | ATR inhibitor | Sigma-Aldrich, SML1415 | 10 µM for 1 h |  |
| NU-7441 | DNA-PK inhibitor | Tocris | 10 µM for 1 h |  |
| DRB | Transcription inhibitor | TCI Deutschland, D4292 | 100 µM for 3 h, as well as during live-cell imaging |  |
| Harmine | DYRK1A inhibitor | Sigma-Aldrich, SMB00461 | 10 µM for 24 h |  |
| INDY | DYRK1A inhibitor | Sigma-Aldrich, SML1011 | 25 µM for 24 h |  |
| **Software** |  |  |  |  |
| Fiji | https://imagej.net/software/fiji/downloads |  |  |  |
| Ilastik-1.4.0 | https://www.ilastik.org/ |  |  |  |
| Graphpad Prism 10 | https://www.graphpad.com/ |  |  |  |
| Adobe Illustrator v25.1 | https://www.adobe.com/de/products/illustrator.html |  |  |  |

<sup>a</sup>Final concentrations in culture medium are indicated

## ACKNOWLEDGEMENTS

We acknowledge the assistance of Fiona Koloßa, Tabea Kleitke and Felix Illner in cloning the open reading frames of hnRNPA2B1, hnRNPR and hnRNPL, respectively. The FOKI U2OS cells were a kind gift from Roger Greenberg (University of Pennsylvania, US) and the 53BP1-MFFR cDNA was a kind gift from Amélie Fradet-Turcotte (Université Laval, Canada). We acknowledge support from the Deutsche Forschungsgemeinschaft (DFG, CRC 1678, grant agreement no. 520471345, as well as FOR 5504, project no. 496650118), the Federation of European Biochemical Societies (FEBS, FEBS Excellence Award). E.V. was supported by the European Union’s Horizon Europe research and innovation programme (Marie Sklodowska-Curie Actions postdoctoral fellowship, grant agreement no. 101150643). Views and opinions expressed are those of the authors only and do not necessarily reflect those of the European Union or the European Research Executive Agency (REA). Neither the European Union nor the granting authority can be held responsible for them. The funders had no role in study design, data collection and analysis, decision to publish, or preparation of the manuscript.

## SUPPLEMENTARY FIGURE LEGENDS

**Supplementary Figure 1** (A-H) HeLa cells transfected with the indicated constructs were subjected to laser microirradiation. Images were obtained at 10 sec (or 30 sec for B) intervals for 15 min, and fluorescence intensities at the damage sites were quantified. (A) Dynamics of GFP (n = 51) and GFP-53BP1^MFFR^ (n = 31) at laser-damaged sites (left), and representative confocal images of GFP-53BP1^MFFR^ before and after laser microirradiation (right). (B) Dynamics of GFP (n = 13) and GFP-hnRNPA2/B1 (n = 38) at laser-damaged sites (left), and representative confocal images of GFP-hnRNPA2/B1 before and after laser microirradiation (right). (C) Dynamics of GFP (n = 32) and GFP-hnRNPF (n = 31) at laser-damaged sites (left), and representative confocal images of GFP-hnRNPF before and after laser microirradiation (right). (D) Dynamics of GFP (n = 21) and GFP-PTBP1 (n = 32) at laser-damaged sites (left), and representative confocal images of GFP-PTBP1 before and after laser microirradiation (right). (E) Dynamics of GFP (n = 42) and GFP-hnRNPQ (n = 32) at laser-damaged sites (left), and representative confocal images of GFP-hnRNPQ before and after laser microirradiation (right). (F, G) GFP-SAF-A-expressing cells segregated into two populations plotted separately. Dynamics of GFP (n = 33) and GFP-SAF-A (n = 35) at laser-damaged sites (left), and representative confocal images of GFP-SAF-A before and after laser microirradiation (right). (F) Cells, in which GFP-SAF-A accumulated transiently at laser-damaged sites before being excluded (65.7 % of cells) (G) Cells in which GFP-SAF-A was excluded without accumulating above the GFP control (34.3 % of cells). (H) Dynamics of GFP (n = 28) and GFP-hnRNPC (n = 36) at laser-damaged sites (left), and representative confocal images of GFP-hnRNPC before and after laser microirradiation (right). Data are shown as mean ± SD from a representative of 3 independent experiments; two-way ANOVA, n.s. non-significant (p ≥ 0.05), ****p < 0.0001. The red arrows indicate the microirradiated area. Scale bars represent 10 µM. Source data are provided as a Source Data file.

**Supplementary Figure 2** (A) Whole-cell extracts from U2OS cells non-irradiated and irradiated (5 Gy) were separated using SDS-PAGE and analysed for hnRNPH1 levels by immunoblotting. Vinculin and CHK2 are used as loading controls. pCHK2 is used as a control for irradiation. (B) Whole-cell extracts from HeLa cells non-irradiated, irradiated (5 Gy), or treated with a combination of 20 μM ATM inhibitor (KU-55933), 10 μM ATR inhibitor (VE-821) and 10 μM DNA-PK inhibitor (NU-7441) for 1 hour prior to irradiation were separated using SDS-PAGE and analysed for γ-H2AX accumulation by immunoblotting. H2AX and Vinculin are used as loading controls. (C) HeLa cells were transfected with GFP-hnRNPH1, and were treated with H_2_O or 10 μM Harmine, or DMSO or 25 μM INDY for 24 hours prior to laser microirradiation. The red arrows indicate the microirradiated area. Scale bars represent 10 µM. (D) Quantification of (C). Data are shown as mean ± SD from 3 independent experiments; GFP-hnRNPH1+DMSO (n=38), GFP-hnRNPH1+H_2_O (n=37), GFP-hnRNPH1+INDY (n=34), GFP-hnRNPH1+Harmine (n=26); two-way ANOVA, n.s. non-significant (p ≥ 0.05), ****p < 0.0001. (E) Whole-cell extracts from HeLa cells untreated or treated with 100 μM transcription inhibitor (DRB) for 4 hours were separated using SDS-PAGE and analysed for pSer5-CTD, pSer2-CTD and pSer7-CTD accumulation by immunoblotting. RPB1-NTD and Tubulin are used loading controls. Source data are provided as a Source Data file.

**Supplementary Figure 3** (A) Whole-cell extracts from HeLa cells transfected with the GFP-hnRNPH1 full-length (FL) or variants were separated using SDS-PAGE and analysed for protein levels by immunoblotting. GAPDH is used as loading control. Source data are provided as a Source Data file.

**Supplementary Figure 4** (A) Whole-cell extracts from U2OS cells untreated or treated with Doxycyclin (Dox) for 24 hours to induce the expression of the different constructs were separated using SDS-PAGE and analysed for protein levels by immunoblotting. GAPDH is used as a loading control. (B) U2OS cells stably expressing the indicated constructs were irradiated (5 Gy), fixed 1 hour after irradiation and processed for MDC1 immunofluorescence. Scale bars represent 5 μM. Dashed lines indicate nucleus outlines (as determined using DAPI staining; not shown). Insets represent 4x magnifications of the indicated fields. (C) Quantification of (B). Data are shown as superplots, the bar represents the median of 3 independent replicates; at least 100 cells per condition were counted; one-way ANOVA, n.s. non-significant (p ≥ 0.05). (D) Whole-cell extracts from U2OS cells expressing the different constructs were separated using SDS-PAGE and analysed for 53BP1 levels by immunoblotting. GAPDH is used as a loading control. Source data are provided as a Source Data file.

**Supplementary Figure 5** (A) U2OS cells stably expressing the indicated constructs were irradiated (5 Gy), fixed at different time-points after irradiation and processed for 53BP1 immunofluorescence. The number of 53BP1 foci per cell was quantified. Data are shown as superplots, the bar represents the median of 3 independent replicates; at least 100 cells per condition were counted; one-way ANOVA, non-significant (p ≥ 0.05), ****p < 0.0001. (B) U2OS cells stably expressing the indicated constructs were irradiated (5 Gy), fixed at different time-points after irradiation and processed for γ-H2AX immunofluorescence. The number of γ-H2AX foci per cell was quantified. Data are shown as superplots, the bar represents the median of 3 independent replicates; at least 100 cells per condition were counted; one-way ANOVA, non-significant (p ≥ 0.05). (C) U2OS cells stably expressing the indicated constructs were irradiated (5 Gy), fixed at different time-points after irradiation and processed for MDC1 immunofluorescence. The number of MDC1 foci per cell was quantified. Data are shown as superplots, the bar represents the median of 3 independent replicates; at least 100 cells per condition were counted; one-way ANOVA, non-significant (p ≥ 0.05). (D) U2OS cells stably expressing the indicated constructs were irradiated (5 Gy), fixed at different time-points after irradiation and processed for RPA32 immunofluorescence. The number of RPA32 foci per cell was quantified. Data are shown as superplots, the bar represents the median of 3 independent replicates; at least 100 cells per condition were counted; one-way ANOVA, non-significant (p ≥ 0.05).

## REFERENCES

1. Tiwari, V. and Wilson, D.M. (2019) DNA damage and associated DNA repair defects in disease and premature aging. Am. J. Hum. Genet., 105, 237–257.

2. Swift, M.L., Rogers, C.M., Kaur, H., et al. (2026) Mechanisms and disease relevance of DNA break repair pathway choice. Nat. Rev. Mol. Cell Biol., 10.1038/s41580-026-01026-3.

3. Aguilar-Quesada, R., Muñoz-Gámez, J.A., Martín-Oliva, D., et al. (2007) Interaction between ATM and PARP-1 in response to DNA damage and sensitization of ATM deficient cells through PARP inhibition. BMC Mol. Biol., 8, 29.

4. Ali, A.A.E., Timinszky, G., Arribas-Bosacoma, R., et al. (2012) The zinc-finger domains of PARP1 cooperate to recognize DNA strand breaks. Nat. Struct. Mol. Biol., 19, 685–692.

5. Haince, J.-F., Kozlov, S., Dawson, V.L., et al. (2007) Ataxia Telangiectasia Mutated (ATM) Signaling Network Is Modulated by a Novel Poly(ADP-ribose)-dependent Pathway in the Early Response to DNA-damaging Agents. J. Biol. Chem., 282, 16441–16453.

6. Langelier, M.-F., Planck, J.L., Roy, S., et al. (2012) Structural Basis for DNA Damage–Dependent Poly(ADP-ribosyl)ation by Human PARP-1. Science, 336, 728–732.

7. Iacovoni, J.S., Caron, P., Lassadi, I., et al. (2010) High-resolution profiling of γH2AX around DNA double strand breaks in the mammalian genome. EMBO J., 29, 1446–1457.

8. Lee, J.-H. and Paull, T.T. (2005) ATM Activation by DNA Double-Strand Breaks Through the Mre11-Rad50-Nbs1 Complex. Science, 308, 551–554.

9. Rogakou, E.P., Boon, C., Redon, C., et al. (1999) Megabase chromatin domains involved in DNA double-strand breaks in vivo. J. Cell Biol., 146, 905–916.

10. Rogakou, E.P., Pilch, D.R., Orr, A.H., et al. (1998) DNA double-stranded breaks induce histone H2AX phosphorylation on serine 139. J. Biol. Chem., 273, 5858–5868.

11. Uziel, T., Lerenthal, Y., Moyal, L., et al. (2003) Requirement of the MRN complex for ATM activation by DNA damage. EMBO J., 22, 5612–5621.

12. Doil, C., Mailand, N., Bekker-Jensen, S., et al. (2009) RNF168 Binds and Amplifies Ubiquitin Conjugates on Damaged Chromosomes to Allow Accumulation of Repair Proteins. Cell, 136, 435–446.

13. Kolas, N.K., Chapman, J.R., Nakada, S., et al. (2007) Orchestration of the DNA-Damage Response by the RNF8 Ubiquitin Ligase. Science, 318, 1637–1640.

14. Mailand, N., Bekker-Jensen, S., Faustrup, H., et al. (2007) RNF8 Ubiquitylates Histones at DNA Double-Strand Breaks and Promotes Assembly of Repair Proteins. Cell, 131, 887–900.

15. Stucki, M., Clapperton, J.A., Mohammad, D., et al. (2005) MDC1 Directly Binds Phosphorylated Histone H2AX to Regulate Cellular Responses to DNA Double-Strand Breaks. Cell, 123, 1213–1226.

16. Wang, B. and Elledge, S.J. (2007) Ubc13/Rnf8 ubiquitin ligases control foci formation of the Rap80/Abraxas/Brca1/Brcc36 complex in response to DNA damage. Proc. Natl. Acad. Sci., 104, 20759–20763.

17. Bunting, S.F., Callén, E., Wong, N., et al. (2010) 53BP1 Inhibits Homologous Recombination in Brca1-Deficient Cells by Blocking Resection of DNA Breaks. Cell, 141, 243–254.

18. Chapman, J.R., Barral, P., Vannier, J.-B., et al. (2013) RIF1 Is Essential for 53BP1-Dependent Nonhomologous End Joining and Suppression of DNA Double-Strand Break Resection. Mol. Cell, 49, 858–871.

19. Escribano-Díaz, C., Orthwein, A., Fradet-Turcotte, A., et al. (2013) A Cell Cycle-Dependent Regulatory Circuit Composed of 53BP1-RIF1 and BRCA1-CtIP Controls DNA Repair Pathway Choice. Mol. Cell, 49, 872–883.

20. Fradet-Turcotte, A., Canny, M.D., Escribano-Díaz, C., et al. (2013) 53BP1 is a reader of the DNA-damage-induced H2A Lys 15 ubiquitin mark. Nature, 499, 50–54.

21. Dev, H., Chiang, T.-W.W., Lescale, C., et al. (2018) Shieldin complex promotes DNA end-joining and counters homologous recombination in BRCA1-null cells. Nat. Cell Biol., 20, 954–965.

22. Di Virgilio, M., Callen, E., Yamane, A., et al. (2013) Rif1 Prevents Resection of DNA Breaks and Promotes Immunoglobulin Class Switching. Science, 339, 711–715.

23. Feng, L., Fong, K.-W., Wang, J., et al. (2013) RIF1 Counteracts BRCA1-mediated End Resection during DNA Repair. J. Biol. Chem., 288, 11135–11143.

24. Findlay, S., Heath, J., Luo, V.M., et al. (2018) SHLD2/FAM35A co-operates with REV7 to coordinate DNA double-strand break repair pathway choice. EMBO J., 37, EMBJ2018100158.

25. Gao, S., Feng, S., Ning, S., et al. (2018) An OB-fold complex controls the repair pathways for DNA double-strand breaks. Nat. Commun., 9, 3925.

26. Gupta, R., Somyajit, K., Narita, T., et al. (2018) DNA Repair Network Analysis Reveals Shieldin as a Key Regulator of NHEJ and PARP Inhibitor Sensitivity. Cell, 173, 972–988.e23.

27. Noordermeer, S.M., Adam, S., Setiaputra, D., et al. (2018) The shieldin complex mediates 53BP1-dependent DNA repair. Nature, 560, 117–121.

28. Panier, S. and Boulton, S.J. (2014) Double-strand break repair: 53BP1 comes into focus. Nat. Rev. Mol. Cell Biol., 15, 7–18.

29. Tomida, J., Takata, K., Bhetawal, S., et al. (2018) FAM35A associates with REV7 and modulates DNA damage responses of normal and BRCA1-defective cells. EMBO J., 37, EMBJ201899543.

30. Zimmermann, M., Lottersberger, F., Buonomo, S.B., et al. (2013) 53BP1 Regulates DSB Repair Using Rif1 to Control 5ʹ End Resection. Science, 339, 700–704.

31. Ceppi, I., Dello Stritto, M.R., Mütze, M., et al. (2024) Mechanism of BRCA1–BARD1 function in DNA end resection and DNA protection. Nature, 634, 492–500.

32. Cruz-García, A., López-Saavedra, A. and Huertas, P. (2014) BRCA1 Accelerates CtIP-Mediated DNA-End Resection. Cell Rep., 9, 451–459.

33. Salunkhe, S., Daley, J.M., Kaur, H., et al. (2024) Promotion of DNA end resection by BRCA1–BARD1 in homologous recombination. Nature, 634, 482–491.

34. Ceccaldi, R. and Cejka, P. (2025) Mechanisms and regulation of DNA end resection in the maintenance of genome stability. Nat. Rev. Mol. Cell Biol., 26, 586–599.

35. Kilic, S., Lezaja, A., Gatti, M., et al. (2019) Phase separation of 53BP1 determines liquid-like behavior of DNA repair compartments. EMBO J., 38, EMBJ2018101379.

36. Piccinno, R., Minneker, V. and Roukos, V. (2019) 53BP1—DNA repair enters a new liquid phase. EMBO J., 38, EMBJ2019102871.

37. Spegg, V. and Altmeyer, M. (2021) Biomolecular condensates at sites of DNA damage: More than just a phase. DNA Repair, 106, 103179.

38. Wang, Y.-L., Zhao, W.-W., Shi, J., et al. (2023) Liquid-liquid phase separation in DNA double-strand breaks repair. Cell Death Dis., 14, 746.

39. Ochs, F., Karemore, G., Miron, E., et al. (2019) Stabilization of chromatin topology safeguards genome integrity. Nature, 574, 571–574.

40. Altmeyer, M., Neelsen, K.J., Teloni, F., et al. (2015) Liquid demixing of intrinsically disordered proteins is seeded by poly(ADP-ribose). Nat. Commun., 6, 8088.

41. Mortusewicz, O., Fouquerel, E., Amé, J.-C., et al. (2011) PARG is recruited to DNA damage sites through poly(ADP-ribose)- and PCNA-dependent mechanisms. Nucleic Acids Res., 39, 5045–5056.

42. He, S., Huang, Z., Liu, Y., et al. (2024) DNA break induces rapid transcription repression mediated by proteasome-dependent RNAPII removal. Cell Rep., 43, 114420.

43. Pankotai, T., Bonhomme, C., Chen, D., et al. (2012) DNAPKcs-dependent arrest of RNA polymerase II transcription in the presence of DNA breaks. Nat. Struct. Mol. Biol., 19, 276–282.

44. Shanbhag, N.M., Rafalska-Metcalf, I.U., Balane-Bolivar, C., et al. (2010) ATM-Dependent Chromatin Changes Silence Transcription In cis to DNA Double-Strand Breaks. Cell, 141, 970–981.

45. Pradel, L., Marnef, A. and Legube, G. (2026) Transcription in check at DNA double-strand breaks. Trends Genet., 42, 937–952.

46. Francia, S., Michelini, F., Saxena, A., et al. (2012) Site-specific DICER and DROSHA RNA products control the DNA-damage response. Nature, 488, 231–235.

47. Michelini, F., Pitchiaya, S., Vitelli, V., et al. (2017) Damage-induced lncRNAs control the DNA damage response through interaction with DDRNAs at individual double-strand breaks. Nat. Cell Biol., 19, 1400–1411.

48. Pessina, F., Giavazzi, F., Yin, Y., et al. (2019) Functional transcription promoters at DNA double-strand breaks mediate RNA-driven phase separation of damage-response factors. Nat. Cell Biol., 21, 1286–1299.

49. Klaric, J.A., Wüst, S. and Panier, S. (2021) New faces of old friends: emerging new roles of RNA-binding proteins in the DNA double-strand break response. Front. Mol. Biosci.

50. Geuens, T., Bouhy, D. and Timmerman, V. (2016) The hnRNP family: insights into their role in health and disease. Hum. Genet., 135, 851–867.

51. Britton, S., Dernoncourt, E., Delteil, C., et al. (2014) DNA damage triggers SAF-A and RNA biogenesis factors exclusion from chromatin coupled to R-loops removal. Nucleic Acids Res., 42, 9047–9062.

52. Polo, S.E., Blackford, A.N., Chapman, J.R., et al. (2012) Regulation of DNA-End Resection by hnRNPU-like Proteins Promotes DNA Double-Strand Break Signaling and Repair. Mol. Cell, 45, 505–516.

53. Adamson, B., Smogorzewska, A., Sigoillot, F.D., et al. (2012) A genome-wide homologous recombination screen identifies the RNA-binding protein RBMX as a component of the DNA-damage response. Nat. Cell Biol., 14, 318–328.

54. Chen, J.-K., Lin, W.-L., Chen, Z., et al. (2018) PARP-1–dependent recruitment of cold-inducible RNA-binding protein promotes double-strand break repair and genome stability. Proc. Natl. Acad. Sci., 115.

55. Mastrocola, A.S., Kim, S.H., Trinh, A.T., et al. (2013) The RNA-binding Protein Fused in Sarcoma (FUS) Functions Downstream of Poly(ADP-ribose) Polymerase (PARP) in Response to DNA Damage. J. Biol. Chem., 288, 24731–24741.

56. Rulten, S.L., Rotheray, A., Green, R.L., et al. (2014) PARP-1 dependent recruitment of the amyotrophic lateral sclerosis-associated protein FUS/TLS to sites of oxidative DNA damage. Nucleic Acids Res., 42, 307–314.

57. Shu, Y., Zhang, J., Zhou, L., et al. (2026) Deacetylated PCBP1 licenses PARP1 activity for DNA damage repair. Mol. Cell, 86, 2703–2721.e8.

58. Varhoshkova, S., Sang, C.C., Antonova, L., et al. (2026) RNA polymerase II drives FUS and EWSR1 exclusion from damaged chromatin. Cell Rep., 45, 117236.

59. Wang, W.-Y., Pan, L., Su, S.C., et al. (2013) Interaction of FUS and HDAC1 regulates DNA damage response and repair in neurons. Nat. Neurosci., 16, 1383–1391.

60. Woods, N.T., Mesquita, R.D., Sweet, M., et al. (2012) Charting the Landscape of Tandem BRCT Domain–Mediated Protein Interactions. Sci. Signal., 5.

61. Chen, F., Xu, W., Tang, M., et al. (2025) hnRNPA2B1 deacetylation by SIRT6 restrains local transcription and safeguards genome stability. Cell Death Differ., 32, 382–396.

62. dos Santos Passos, C., Choi, Y.-S., Snow, C.D., et al. (2021) Design of genetically encoded sensors to detect nucleosome ubiquitination in live cells. J. Cell Biol., 220, e201911130.

63. Saquilabon Cruz, G.M., Kong, X., Silva, B.A., et al. (2016) Femtosecond near-infrared laser microirradiation reveals a crucial role for PARP signaling on factor assemblies at DNA damage sites. Nucleic Acids Res., 44, e27.

64. Patel, A., Lee, H.O., Jawerth, L., et al. (2015) A Liquid-to-Solid Phase Transition of the ALS Protein FUS Accelerated by Disease Mutation. Cell, 162, 1066–1077.

65. Sun, L., Ye, R., Cao, C., et al. (2024) BCAS2 and hnRNPH1 orchestrate alternative splicing for DNA double-strand break repair and synapsis in meiotic prophase I. Cell. Mol. Life Sci., 81, 449.

66. Wen, C., Tian, Z., Li, L., et al. (2022) SRSF3 and HNRNPH1 Regulate Radiation-Induced Alternative Splicing of Protein Arginine Methyltransferase 5 in Hepatocellular Carcinoma. Int. J. Mol. Sci., 23, 14832.

67. Gall-Duncan, T., Luo, J., Jurkovic, C.-M., et al. (2023) Antagonistic roles of canonical and Alternative-RPA in disease-associated tandem CAG repeat instability. Cell, 186, 4898–4919.e25.

68. Maréchal, A., Li, J.-M., Ji, X.Y., et al. (2014) PRP19 Transforms into a Sensor of RPA-ssDNA after DNA Damage and Drives ATR Activation via a Ubiquitin-Mediated Circuitry. Mol. Cell, 53, 235–246.

69. Kong, X., Mohanty, S.K., Stephens, J., et al. (2009) Comparative analysis of different laser systems to study cellular responses to DNA damage in mammalian cells. Nucleic Acids Res., 37, e68–e68.

70. Guard, S.E., Poss, Z.C., Ebmeier, C.C., et al. (2019) The nuclear interactome of DYRK1A reveals a functional role in DNA damage repair. Sci. Rep., 9, 6539.

71. R. Menon, V., Ananthapadmanabhan, V., Swanson, S., et al. (2019) DYRK1A regulates the recruitment of 53BP1 to the sites of DNA damage in part through interaction with RNF169. Cell Cycle, 18, 531–551.

72. Roewenstrunk, J., Di Vona, C., Chen, J., et al. (2019) A comprehensive proteomics-based interaction screen that links DYRK1A to RNF169 and to the DNA damage response. Sci. Rep., 9, 6014.

73. Kim, G.H. and Kwon, I. (2021) Distinct roles of hnRNPH1 low-complexity domains in splicing and transcription. Proc. Natl. Acad. Sci., 118, e2109668118.

74. Huen, M.S.Y., Grant, R., Manke, I., et al. (2007) RNF8 Transduces the DNA-Damage Signal via Histone Ubiquitylation and Checkpoint Protein Assembly. Cell, 131, 901–914.

75. Mirman, Z., Sasi, N.K., King, A., et al. (2022) 53BP1–shieldin-dependent DSB processing in BRCA1-deficient cells requires CST–Polα–primase fill-in synthesis. Nat. Cell Biol., 24, 51–61.

76. Swift, M.L., Zhou, R., Syed, A., et al. (2023) Dynamics of the DYNLL1–MRE11 complex regulate DNA end resection and recruitment of Shieldin to DSBs. Nat. Struct. Mol. Biol., 30, 1456–1467.

77. Harrigan, J.A., Belotserkovskaya, R., Coates, J., et al. (2011) Replication stress induces 53BP1-containing OPT domains in G1 cells. J. Cell Biol., 193, 97–108.

78. Lukas, C., Savic, V., Bekker-Jensen, S., et al. (2011) 53BP1 nuclear bodies form around DNA lesions generated by mitotic transmission of chromosomes under replication stress. Nat. Cell Biol., 13, 243–253.

79. Chen, R. and Wold, M.S. (2014) Replication protein A: Single-stranded DNA’s first responder: Dynamic DNA-interactions allow replication protein A to direct single-strand DNA intermediates into different pathways for synthesis or repair. BioEssays, 36, 1156–1161.

80. Banani, S.F., Lee, H.O., Hyman, A.A., et al. (2017) Biomolecular condensates: organizers of cellular biochemistry. Nat. Rev. Mol. Cell Biol., 18, 285–298.

81. Kilgas, S., Syed, A., Toolan-Kerr, P., et al. (2024) NEAT1 modulates the TIRR/53BP1 complex to maintain genome integrity. Nat. Commun., 15, 8438.

82. Tanu, T., Taniue, K., Imamura, K., et al. (2021) hnRNPH1-MTR4 complex-mediated regulation of NEAT1v2 stability is critical for IL8 expression. RNA Biol., 18, 537–547.

83. Drané, P., Brault, M.-E., Cui, G., et al. (2017) TIRR regulates 53BP1 by masking its histone methyl-lysine binding function. Nature, 543, 211–216.

