## Supplementary Figures for "Exclusion from DNA double-strand breaks is a pervasive feature of the hnRNP family"

Supplementary Figure 1 Vincendeau et al.

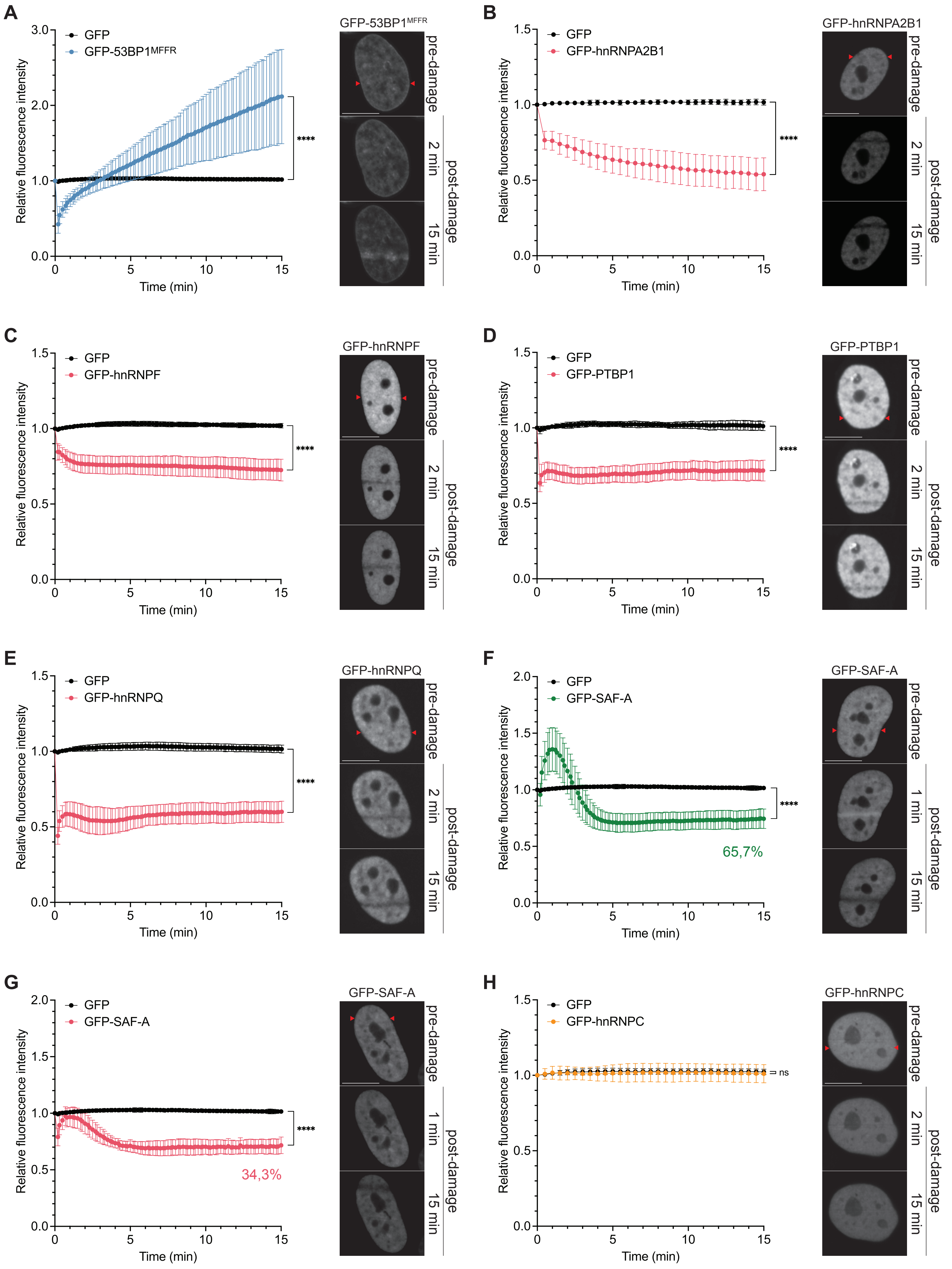

Supplementary Figure 2 Vincendeau et al.

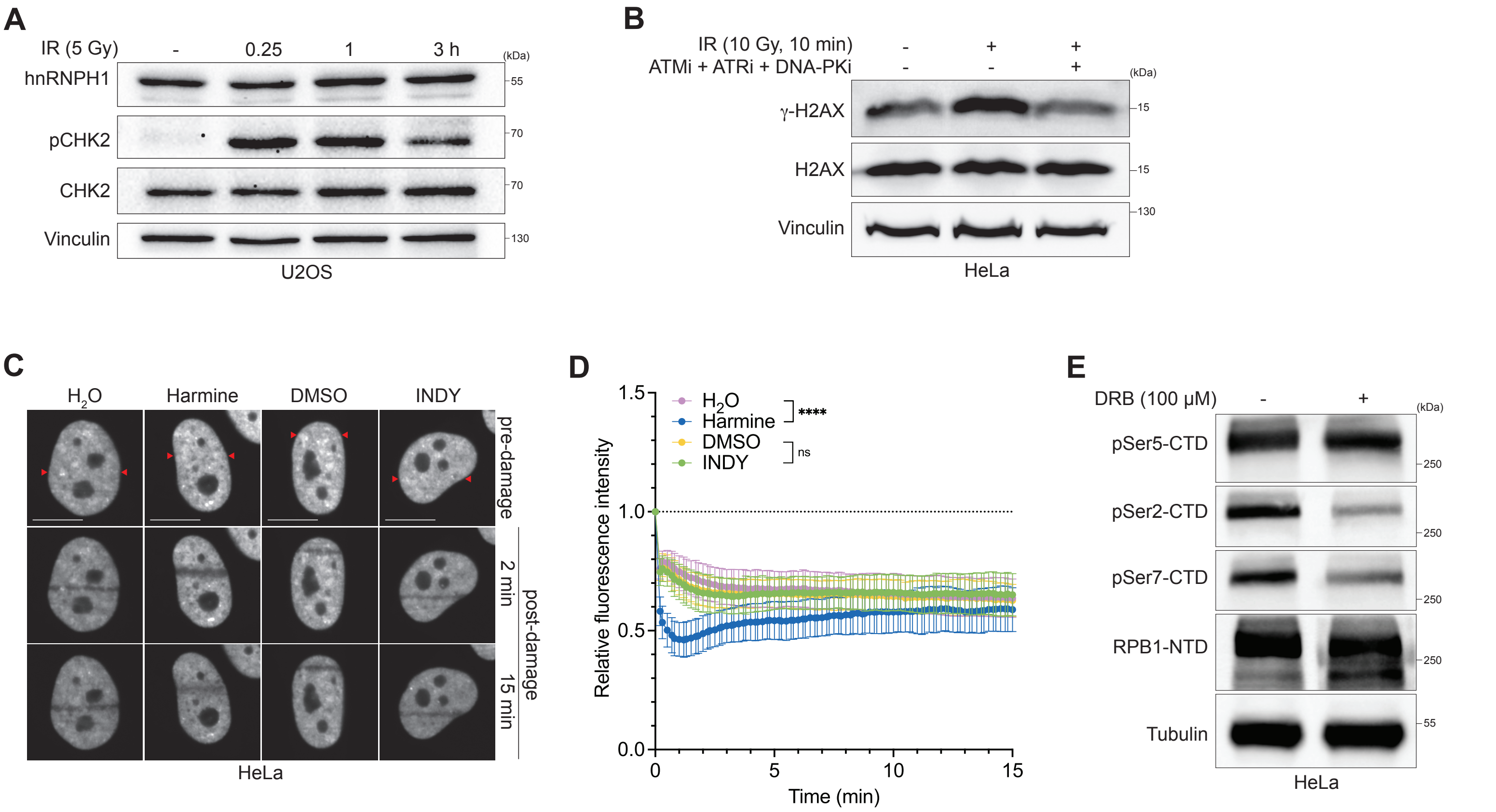

**A**

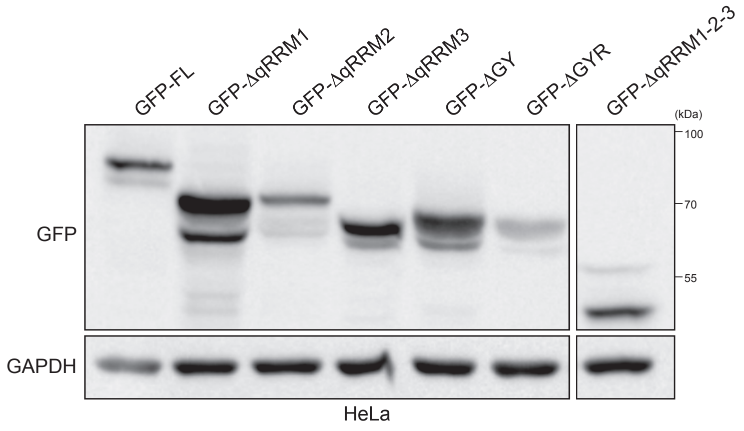
GFP-FL
GFP-ΔqRRM1
GFP-ΔqRRM2
GFP-ΔqRRM3
GFP-ΔGY
GFP-ΔGYR
GFP-ΔqRRM1-2-3
(kDa)
100
70
55
GFP
GAPDH
HeLa

Supplementary Figure 4 Vincendeau et al.

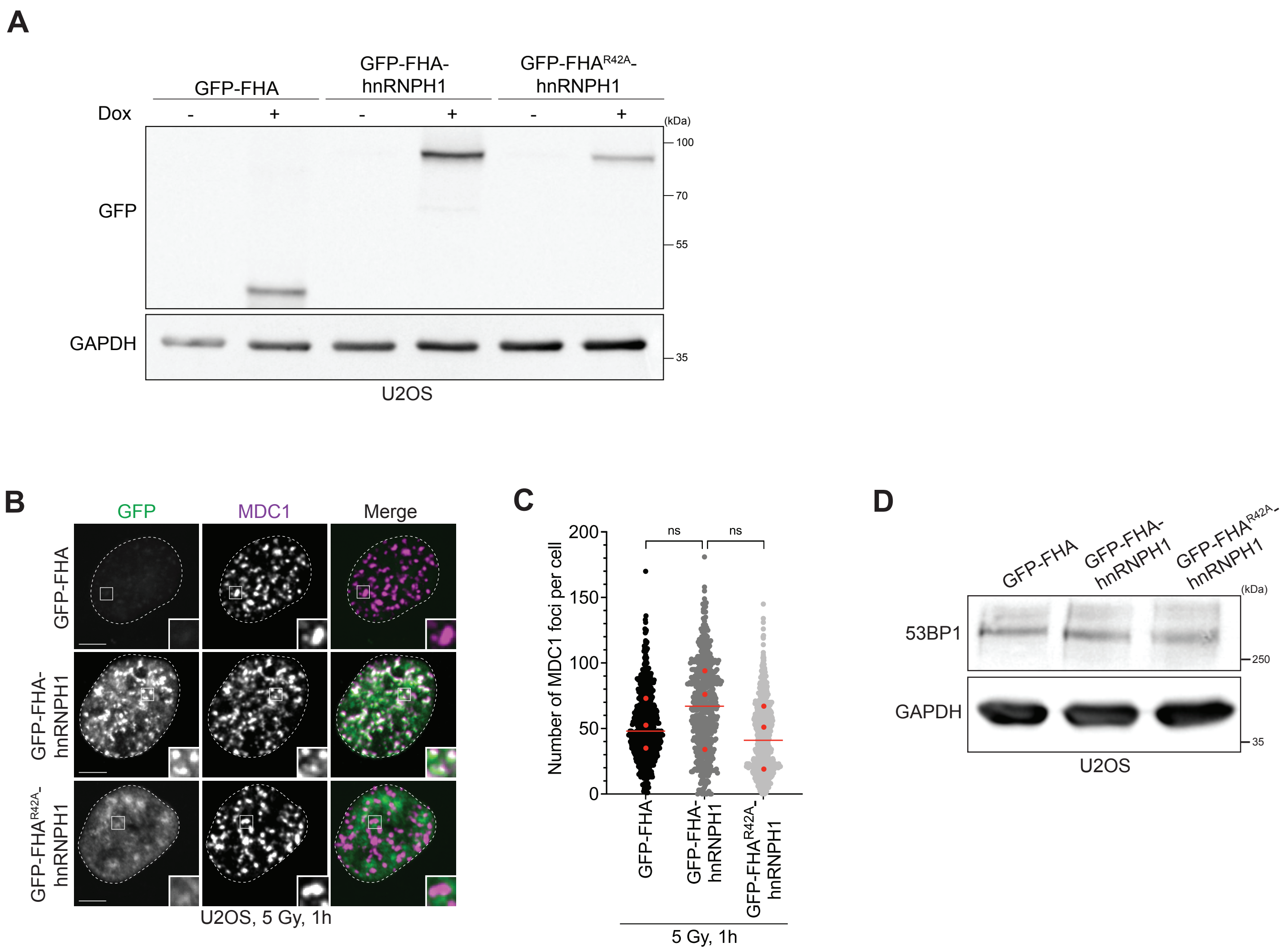

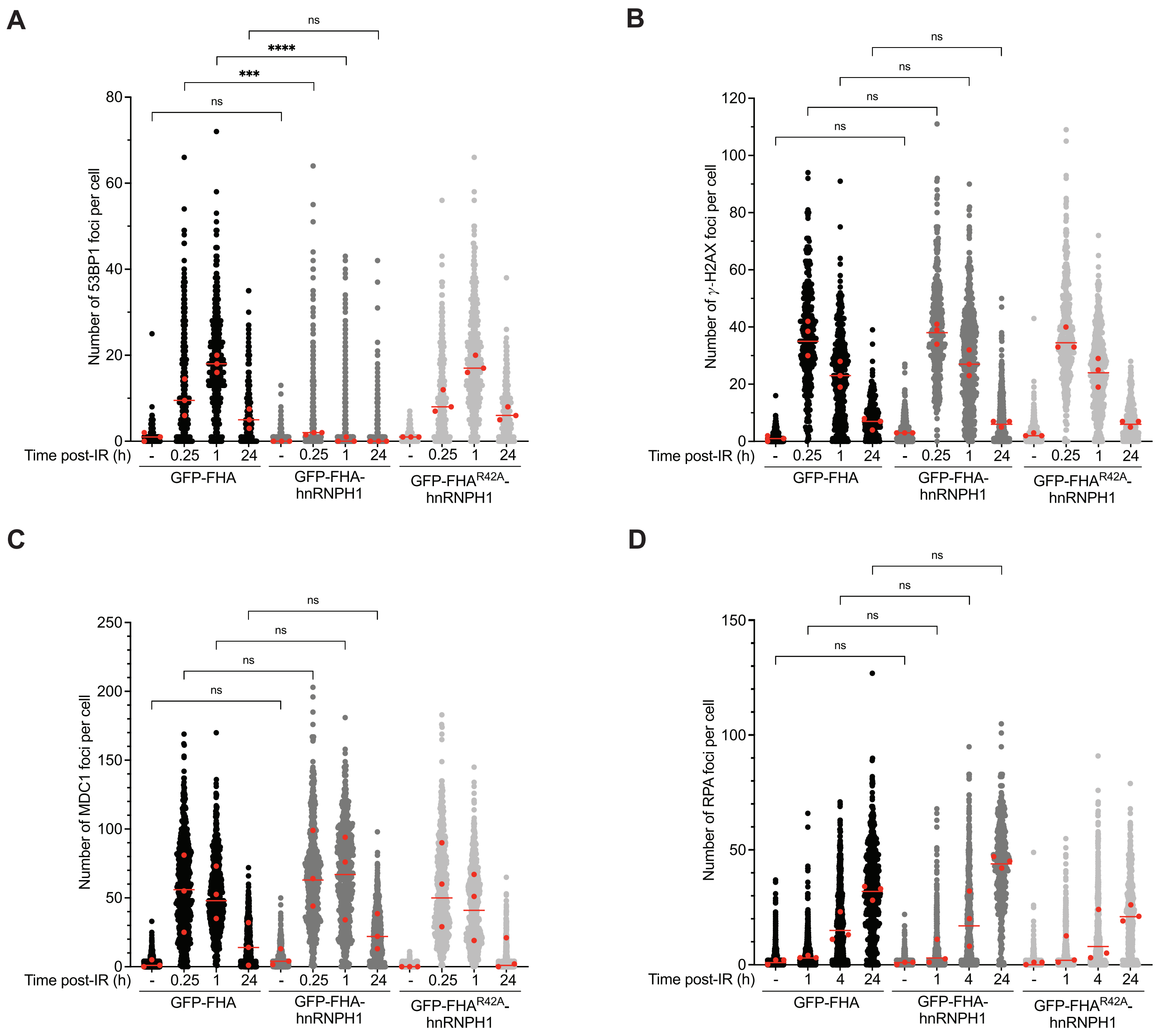
A
Number of 53BP1 foci per cell
ns
****
***
ns
Time post-IR (h)
- 0.25 1 24
GFP-FHA
GFP-FHA-hnRNPH1
GFP-FHA$^{R42A}$-hnRNPH1
B
Number of γ-H2AX foci per cell
ns
ns
ns
ns
Time post-IR (h)
- 0.25 1 24
GFP-FHA
GFP-FHA-hnRNPH1
GFP-FHA$^{R42A}$-hnRNPH1
C
Number of MDC1 foci per cell
ns
ns
ns
ns
Time post-IR (h)
- 0.25 1 24
GFP-FHA
GFP-FHA-hnRNPH1
GFP-FHA$^{R42A}$-hnRNPH1
D
Number of RPA foci per cell
ns
ns
ns
ns
Time post-IR (h)
- 1 4 24
GFP-FHA
GFP-FHA-hnRNPH1
GFP-FHA$^{R42A}$-hnRNPH1
